# Lymphoma response to chimeric antigen receptor T cell therapy is driven by endogenous tumor-reactive CD8⁺ T cells

**DOI:** 10.64898/2026.10.23.761341

**Authors:** Bachisio Ziccheddu, Constanza Savid-Frontera, Venu V.G. Saralamma, Chandler Gatenbee, Abdessamad A. Youssfi, Monika Chojnacka, Meghan Menges, Luis Cuadrado-Delgado, Michael Durante, John Mullinax, Kayla Reid, Julieta Abraham Miranda, Salvatore Corallo, Emily Merritt, William Sawyer, Anthony Augello, Austin D. Newsam, Dhanvantri Chahar, Catalina Amador, Jay Y. Spiegel, Marco Davila, Ola Landgren, Juan Pablo Alderuccio, Mark Robertson-Tessi, Reginald Atkins, Saad Usmani, Michael D. Jain, Alexander R.A. Anderson, Francesco Maura, Jonathan H. Schatz, Frederick L. Locke

## Abstract

CD19-directed chimeric antigen receptor T-cell (CAR-T) therapy induces durable remissions in a subset of patients with relapsed or refractory large B-cell lymphoma; however, most patients fail to respond or eventually relapse. By integrating whole-genome and transcriptome profiling of 54 patients with functional assays, syngeneic models, and mathematical modeling, we demonstrate that durable CAR-T efficacy requires cooperation between CAR-T cells and endogenous CD8⁺ (eCD8) T cells recruited to join the attack on malignant cells. Tumors with biallelic loss of MHC class I or *CD58* exhibited high genomic complexity, increased neoantigen burden, and resistance to CAR-T therapy. MHC-I–deficient lymphomas showed suboptimal response in immunocompetent hosts compared to wild type. Importantly, sustained responses in mice occurred only in the presence of eCD8 cells, whereas all CD8⁺-depleted mice relapsed after initial CAR-T-mediated tumor regression. Patients and murine ELISPOT assays confirmed enhanced endogenous T-cell activation post-CAR-T. These findings reveal that CAR-T therapy functions not only as a cytotoxic modality but also as a catalyst for endogenous anti-tumor immunity.

## INTRODUCTION

CD19-targeted chimeric antigen receptor T-cell therapy (CAR-T) is a transformative treatment for relapsed/refractory large B-cell lymphoma (r/r LBCL), leading to complete and durable responses in 30-40% of patients. However, the remainder unfortunately will relapse or fail to respond, and the mechanisms underlying treatment failures remain incompletely understood^1,2^.

Multiple resistance mechanisms have been described, including permanent loss or reduction of the target antigen, limited CAR T-cell expansion, and immunosuppressive tumor microenvironments^3,4^. In our previous studies, we independently identified two distinct molecular contributors to treatment failure: first, tumor microenvironmental features associated with poor response^5^; and second, tumor-intrinsic genomic alterations^6^. The latter findings align with independent studies demonstrating that tumor genomic complexity, as measured by copy number alterations found within circulating cell-free DNA, is predictive of refractory disease and suboptimal response to anti-CD19 CAR-T ^3,7,8^. Nevertheless, it is unclear how genomic complexity promotes refractoriness and suboptimal LBCL response to CAR-T.

Although CAR-T cells were originally designed to directly eliminate tumor cells expressing the target antigens, emerging evidence suggests that their clinical efficacy in lymphoma may also be dependent on additional factors. For instance, we identified a patient harboring a subclonal CD19 mutation (L124V, previously associated with resistance to CAR-T) who nonetheless experienced a prolonged remission^9^. As additional evidence, a recent report demonstrated that clonal expansion of the native cytotoxic T-cell repertoire correlates with clinical response^10^, supporting the hypothesis that anti-CD19 CAR-T efficacy may, in part, be mediated by CAR T-independent mechanisms^11^. Moreover, preclinical glioblastoma models and patient samples further demonstrate that the host immune system plays a central role in sustaining CAR T-cell anti-tumor activity by reinvigorating host immunity within an otherwise exhausted tumor microenvironment^12^. Thus, CAR T cells may function not only as direct cytotoxic effectors but also as facilitators of endogenous immune activation against tumor cells^13^. Collectively, these observations highlight the need to better understand the dynamic interplay between CAR T cells, tumor cells, and the host immune system, and how this ecosystem shapes therapeutic efficacy and resistance.

To investigate the role of host immunity in CAR-T, we interrogated 61 whole-genome sequencing (WGS) and 54 RNA sequencing (RNAseq) samples from 54 LBCL patients treated with anti-CD19 CAR-T. Our analysis revealed that increased genomic complexity, and biallelic loss of MHC class I (MHC-I) were all associated with worse clinical outcomes following CAR T cell infusion. Complementing these findings, preclinical and ex vivo experiments demonstrated that non-CAR eCD8+ T cells contribute to LBCL tumor clearance after CAR-T infusion. These results suggest that effective tumor eradication requires both sustained CD19 expression and retained immune synapse formation by eCD8 cells with malignant lymphocytes. We propose that CAR T-cell therapy not only directly targets tumor cells but also acts as a catalyst for host immune activation. Moreover, escape from non-CAR T-cells is a novel and significant driver of CAR-T resistance.

## RESULTS

### Patient cohort

A total of 61 tumor biopsies (with paired germline samples) from 54 patients with r/r LBCL treated with CD19-directed CAR T-cell therapy (CD19 CAR-T) were included in the study. Thirty-nine samples were obtained at baseline prior to lymphodepletion and CAR T infusion, eight were collected at disease relapse, and seven patients contributed paired samples before and after treatment (**Fig. 1a**). Sixty-one and fifty-four samples were investigated by WGS and RNAseq, respectively (**Supplementary Table 1**). Demographics, disease characteristics, toxicities, and response to CD19 CAR-T for all patients are summarized in **Supplementary Table 2**. Eight patients had relapse-only biopsies: 2 refractories (CAR_26 and CAR_218), 2 with minimal response (CAR_ 237 and Seq_4), and 4 with transient complete remission (CAR_198, CAR_43, CAR_196 and Seq_3). In the first four, refractoriness or limited response likely reflected preexisting clonal alterations, and they were retained in survival analysis. In the remaining four, complete remission followed by progression may reflect selection of a subclone genomically distinct from the pretreatment tumor, and they were excluded from progression-free survival (PFS) and overall survival (OS) analyses (**Supplementary Fig.1**; **Supplementary Methods**). One patient died within a week after infusion because of CAR-19 toxicity with unknown disease response (CAR_122). This patient was omitted from PFS but included in OS analyses (**Supplementary Fig.1**; **Supplementary Methods**). Median PFS and OS for the entire cohort were 7.85 and 16.2 months, respectively (**Fig. 1b-d)**. Patients were categorized as “durable responders” (complete or partial response sustained to at least 9 months) or “non-durable responders” (no response or relapsed disease). Patients were defined as refractory if they never achieved partial response (primary refractory). Durable response was seen in 18 (33.3%) patients after median post treatment follow up of 24 months (**Supplementary Table 2**). Overall, results are comparable to previously reported CD19 CAR-T outcomes^1,2,14,15^. Investigating features associated with poor outcome after CD19 CAR-T, sex, age and prior lines of treatment associated with high risk of relapse (**Supplementary Table 2**).

**Fig. 1.**
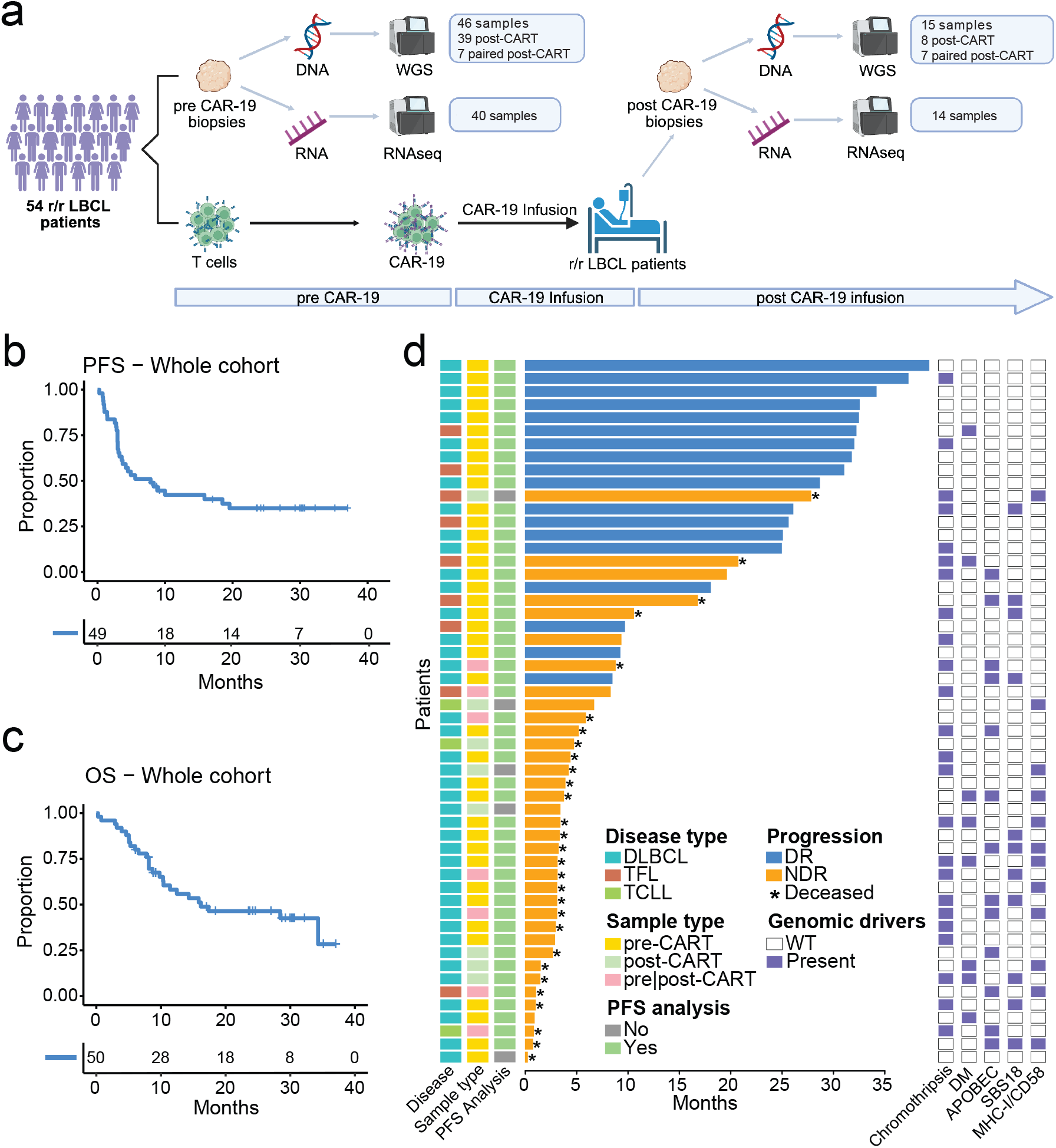
Clinical outcomes and genomic features in r/r LBCL treated with CD19 CAR T cells. **a.** Study design. **b.** PFS Kaplan-Meier curve for the entire cohort. **c.** OS Kaplan– Meier curve for the entire cohort. **d.** Swimmer patient-level plot summarizing the clinical outcome of 54 patients with r/r LBCL. Durable responders (DR) are shown in bleu and non-durable responders (NDR) in orange. The asterisk indicates the death of the patient. The right panel depicts significant genomic features and alterations involving the immune synapse (*HLA-A*, *HLA-B*, *HLA-C*, *B2M* and *CD58*).

### Genomic features and complexity

Expanding upon our previous study^6^, APOBEC activity (SBS2/13), oxidative damage-related mutations (SBS18), chromothripsis, and double minutes, were again associated with inferior progression-free survival (PFS) after CAR-19 therapy (**Supplementary Fig.2 a-g**; **Supplementary Table 3**). As these features largely reflect high levels of genomic instability and genomic complexity, we next leveraged copy number variation (CNV) and structural variant (SV) signatures to infer genomic complexity in DLBCL, as previously done in solid tumors (**Supplementary Fig.3 a-c**; **Supplementary Table 4-5**)^16,17^. Patients harboring a complex genome signature experienced significantly shorter PFS after CD19 CAR-T treatment (p = 0.031, log-rank test; **Fig. 2a**). Collectively, these results confirm that pre-existing genomic complexity is a major determinant of poor outcomes following CD19 CAR-T therapy, consistent with prior observations^3,6,7^.

**Fig. 2.**
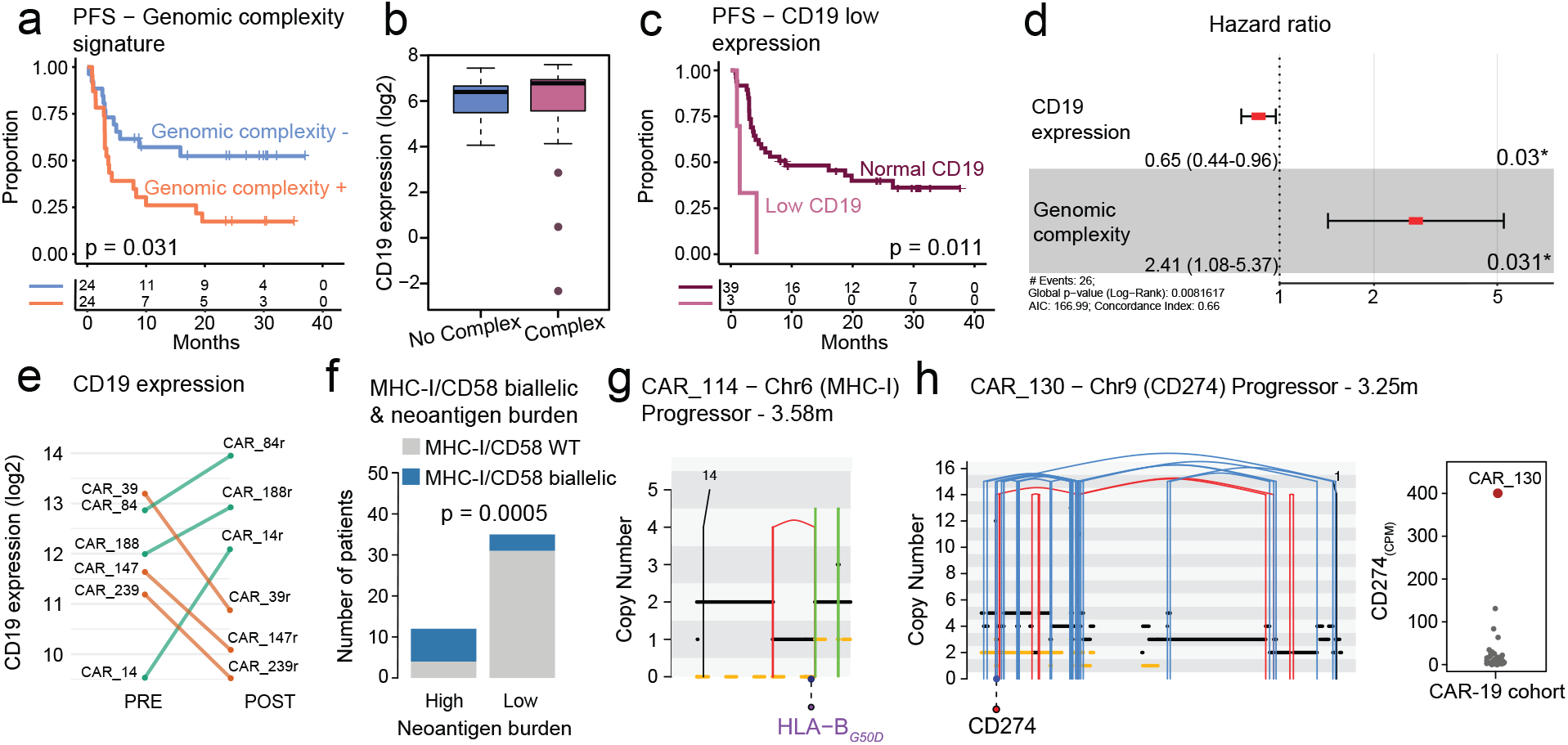
Tumor-intrinsic genomic complexity, CD19 expression, and antigen-presentation defects shape CD19 CAR-T outcome in r/r LBCL. **a.** PFS Kaplan–Meier curve stratified by the complex-genome signature (Signature– vs Signature+); p-value computed by log-rank test. **b.** Boxplot showing the *CD19* expression in patients with and without genomic complexity. **c.** Kaplan-Meir curve of PFS stratified by baseline *CD19* expression in patients with available pretreatment RNAseq data who met the PFS analysis inclusion criteria (N=42); p-value computed by log-rank test. **d.** Multivariate cox model evaluating the hazard ratio (HR) for *CD19* expression and genomic complexity; the ribbon indicates the pointwise 95% confidence interval (HR < 1 denotes lower risk with higher *CD19*). e. *CD19* expression for paired samples. PRE = pre-CAR, POST = post-CAR. Lines connect samples from the same patient; green and orange indicate increased and decreased *CD19* expression after therapy, respectively. **f.** Bar plot showing enrichment of neoantigen burden in patients with biallelic inactivation of immune-synapse (IS) genes (*HLA-A*, *HLA-B*, *HLA-C*, *B2M* and *CD58*); in blue patients harboring MHC-I/CD58 biallelic inactivation, in grey patients with MHC-I/CD58 WT; p-value computed using Fisher’s Exact test (WT: wild type). **g**. Representative chromosome-6 copy-number profiles illustrating biallelic MHC-I disruption in a progressor (CAR_114). Vertical lines: Red = deletion, blue = inversion, green = duplications, black = translocation. Gene names: violet = mutated, red = WT. **h.** Left: CNVs/SVs profile of chromosome 9 in CAR_130 showing chromothripsis involving *CD274*. Right: boxplot displaying the *CD274* gene expression in our cohort, CAR_130 is highlighted in red.

Although permanent loss of *CD19* is an uncommon resistance mechanism, multiple studies have shown that low *CD19* protein and gene expression predict poor outcomes following CAR T cell therapy^18–21^. Analysis of RNA-seq data revealed three patients with marked downregulation of *CD19* expression (6.8%), all of whom exhibited complex genomic profiles and rapid disease progression (p=0.011, log-rank test; **Fig. 2b,c**). No genomic alterations in *CD19* were identified in these three patients. In contrast, in addition to the previously reported case with a subclonal *CD19* mutation (CAR_88), we identified another patient (CAR_64) harboring a *CD19* G210D mutation on a diploid background, a variant previously linked to impaired cell-surface localization of *CD19*^9^. Notably, like the earlier case, this patient showed preserved *CD19* expression by both RNA sequencing and immunohistochemistry and achieved a durable response to CAR-T therapy. In multivariate analysis, both *CD19* downregulation and genomic complexity independently and additively contributed to the risk of progression after CAR T-cell therapy (**Fig. 2d**). Collectively, these findings underscore the major impact of tumor-intrinsic genomic and transcriptomic features on post-CAR T-cell efficacy.

### Clonal evolution post CD19 CAR-T treatment

Across the seven patients with longitudinal sampling before CAR-T and at progression, we detected no genomic events involving *CD19*, and *CD19* transcript levels did not change significantly (three patients downregulated, three upregulated; **Fig. 2e and Supplementary Fig. 4**). In the broader, non-paired cohort, *CD19* expression was likewise comparable between pre- and post-CAR-T specimens (p=0.76; Wilcoxon test). Phylogenetic reconstruction in these seven patients showed limited post-treatment evolution: two tumors followed a linear and five a branching pattern (**Supplementary Fig. 4a-g**), with a dominant clonal shift in three cases (42.9%). Alterations were consistent across anatomical sites, indicating global tumor hallmarks rather than site-specific subclones. In every case the trunk was dominated by chromothripsis, biallelic *TP53* inactivation, APOBEC mutagenesis, and hallmark *MYC*/*BCL2*/*BCL6* lesions; in two patients (CAR_14 and CAR_147; **Supplementary Fig. 4f-g**) the trunk also carried an immune-evasion lesion (biallelic *CD58* inactivation), and both progressed within three months of infusion. In CAR_39, a clonal HLA-B missense variant with predicted functional impact was detected exclusively on the progression branch (**Supplementary Fig. 4e**). Together with the truncal biallelic CD58 inactivation in CAR_14 and CAR_147, it suggests that impairment of T-cell–mediated recognition recurs as a route to CAR-T failure. Among tumors that progressed with a new dominant clone, events selected at progression involved NF-κB signaling (*TNFAIP3*, *NFKBIE*), transcriptional regulators (*FOXO1*, *MEF2B*, *EP300*, *ETV6*), and focal gains (*CCND3*, *NFKBIE*) or *MYC* duplication/gain. Together, these data indicate that the molecular determinants of CD19 CAR-T failure are largely present at the time of infusion and persist through relapse, with only limited post-treatment diversification.

### Tumor immunogenicity is shaped by MHC-I integrity and neoantigen burden

Data from this and other studies demonstrated a strong association between genomic complexity and clinical outcomes following CAR T-cell therapy. In certain solid tumors, genomic complexity and instability have been linked to a higher neoantigen burden and increased responsiveness to immunotherapy (e.g., mismatch repair-deficient cancers)^22,23^. Conversely, genomic complexity might also drive the emergence of immune escape mechanisms. To assess this, we integrated WGS and RNAseq data to predict clonal neoantigens from expressed somatic mutations while accounting for each patient’s specific MHC-I genotype and antigen presentation capacity (n= 54; **Supplementary Fig. 5**; **Supplementary Methods**). The total number of predicted clonal neoantigens did not significantly differ between durable responders and non-durable responders (**Supplementary Fig. 6a**), nor between complex and simple (**Supplementary Fig. 6b**). However, non-durable responders showed a distinctly bimodal distribution of neoantigen burden (**Supplementary Fig. 6a**). Interestingly, patients with high neoantigen burden (n=12; number of neoantigen ≥ 15) were enriched for biallelic loss of genes involved *MHC-I/CD58* compared to the other patients (p=0.0005, Fisher’s exact test; **Fig. 2f**). Furthermore, all but one patient with high neoantigen burden showed evidence of genomic complexity and/or biallelic loss of *MHC-I/CD58* (**Supplementary Table 3**); the one genomically simple patient (CAR_276) was the only one who experienced a prolonged response. These findings drove us to investigate more broadly the impact of genes related to the immune synapse, and genes that play a role in interactions with T-cell effector function. We identified 10 patients with biallelic inactivation of *MHC-I* (7 pre-CAR-T; **Fig. 2g**; **Supplementary Fig. 7a-i; Supplemental Table 3,6**), and 5 with biallelic inactivation of *CD58* (all pre-CART, 2 patients with also biallelic inactivation of *MHC-I*; **Supplementary Fig. 7j-n; Supplemental Table 3,7**). Furthermore, we found one refractory patient (CAR_130) harboring a chromothripsis generating a focal (<5Mb) high level copy-number amplification (12 copies) of *CD274*, causing its upregulation (**Fig. 2h**). Interestingly, all patients with permanent loss of *MHC-I* or *CD58* (**Fig. 3a**) were refractory to CAR-T (p<0.0001, log-rank test; **Fig. 3b**). Conversely, monoallelic loss of *MHC-I* or *CD58*, identified in 21 patients (38.9%) and 12 patients (22.2%), respectively, did not associate with outcome. Importantly *MHC-I/CD58* loss remained independently associated with progression in multivariate cox model (HR 5.73, p=0.008; **Fig 3c**).

**Fig. 3.**
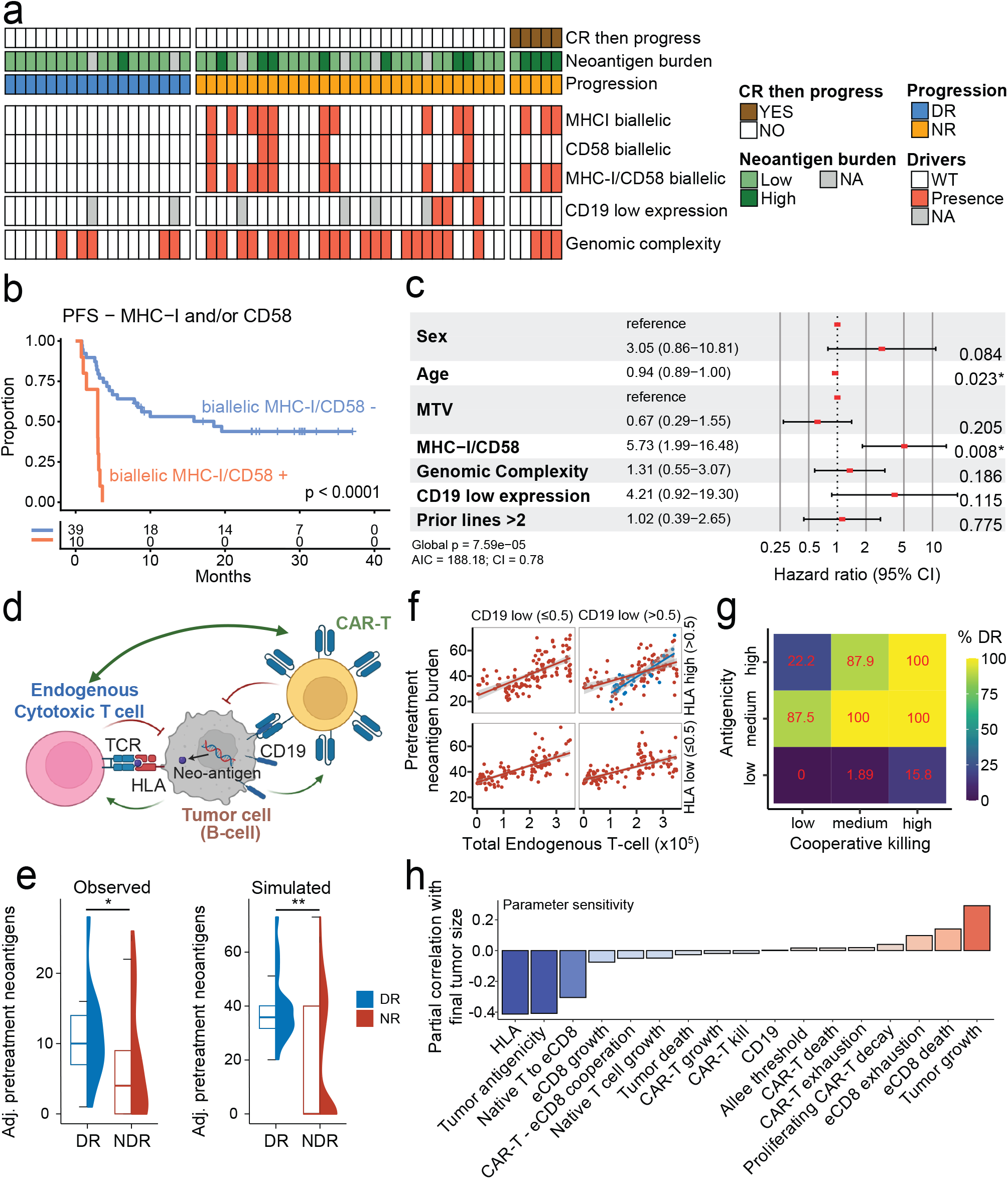
Biallelic disruption of the immune synapse independently predicts CD19 CAR-T failure in r/r LBCL, and a multi-predator model supports cooperation between CAR-T and endogenous CD8 T cells as a requirement for durable response. **a.** Heatmap representing the CAR-19 cohort and the events involving MHC-I/CD58 and genomic complexity (DR: Durable Responders, NDR: Non-Durable Responders; CR: complete remission; WT: wild type). **b.** PFS Kaplan-Meier curve stratified by biallelic MHC-I loss and/or biallelic CD58 loss; p-value computed by log-rank test. **c.** multivariate cox model including sex, age metabolic tumor volume (MTV), genomic complexity, and biallelic MHC-I/CD58 inactivation. **d.** Conceptual model: CD19 CAR-T efficacy engages both direct CD19-mediated tumor recognition and endogenous CD8 T-cell recognition of MHC-I–presented neoantigens. **e.** Relationship between adjusted pretreatment neoantigen burden (based on immune-evasion mechanisms) and progression (observed-based analysis, p = 0.02; simulation-based analysis, p = 0.002). **f.** Simulated pre-treatment neoantigen burden as a function of intra-tumor endogenous CD8 infiltration, stratified by CD19 and HLA expression (low <= 0.5, high > 0.5, with a range of 0-1). **g.** Percentage of simulations achieving a durable response (%DR) across levels of tumor antigenicity and eCD8/CAR-T cooperative killing. **h.** Sensitivity analysis to measure the impact of each model parameter/component on final tumor size. Negative (blue) values indicate that increasing the parameter reduces tumor size, while positive (red) values indicate that increasing the parameter increases tumor size.

Since CAR-T cells engage and eliminate tumor cells independently of MHC-I recognition, these findings raise the possibility that non-CAR endogenous CD8⁺ T cells contribute to durable remissions.

### Computational modeling supports endogenous and CAR-T cooperation

To assess the role of endogenous T and CAR-T cell cooperation, and whether permanent loss of immune synapse-related genes is sufficient to drive resistance to CD19 CAR-T therapy, we developed a multi-predator (endogenous T and CAR) multi-prey (tumor clones) consumer-resource model^24^, in which CAR-T and intra-tumor endogenous CD8 T cells (eCD8) interact through cooperative killing^25^, whereby higher predator abundance increases tumor detection and killing (**Fig. 3d**; **Methods**). The biological rationale is that both T cell populations release pro-inflammatory cytokines that increase activity and chemokines that recruit eCD8 cells to their targets. MHC-I expression on tumor cells determines how readily those cells are detected by eCD8, while CD19 levels determine the CAR-T detection rate. This model is intended to explore the biological plausibility of CAR-T/eCD8 cooperation rather than to provide quantitative patient-level predictions. Critically, model parameters were set from an independent dataset^26^, and the simulated neoantigen comparison draws on a cohort separate from the one used for parameterization, so concordance with our genomic data is not a product of fitting the model to those data. With these independent parameters, the model recapitulated our findings: simulated tumors with both high CD19 and high MHC-I expression demonstrated effective elimination, while those deficient in MHC-I or neoantigen presentation were more likely to progress (**Fig. 3e**). Notably, neoantigen burden was a strong predictor of intra-tumor eCD8 infiltration and treatment success only in the context of intact MHC-I expression (**Fig. 3f**). Simulated tumors with high CD19 but low HLA expression failed to recruit eCD8 cells and were incompletely cleared by CD19 CAR-T alone, whereas those with low CD19 but high MHC-I did not trigger sufficient CAR-mediated cytotoxicity, indicating that both recognition pathways are needed for a durable response. Simulating a range of eCD8/CAR-T cooperation showed that the proportion of durable responses rose with the degree of cooperative killing, but only in tumors of sufficient antigenicity; at low antigenicity, durable responses remained rare even at the highest cooperation simulated (**Fig. 3g**). We then examined the contribution of each parameter to final tumor size (**Fig. 3h**), which serve as a surrogate measure for the balance between tumor growth and immune control, smaller residual tumor burden indicating a greater likelihood of durable response. Because LBCL outcomes after CAR-T therapy are largely dichotomous, final simulated tumor size serves as a surrogate for the balance between tumor growth and immune control, smaller residual burden indicating a greater likelihood of durable response. Unsurprisingly, HLA expression, the immune kill rate, and conversion from naive T cells to eCD8 had the strongest impact on reducing tumor size, as did the cooperative killing parameter (CAR-T/eCD8 interaction in **Fig. 3h**). Parameters related to eCD8 tend to be more impactful than those related to CAR-T, suggesting that CAR-T cells reduce tumor burden not only by direct killing but by amplifying the eCD8 response. This model reinforces the biological synergy observed in our genomic data and supports escape from non-CAR eCD8 cells as a driver of CAR-T resistance, and that cooperation between eCD8 and CAR-T is critical to durable response. We next tested whether immune synapse loss is itself required for resistance.

### CD19 CAR-T success depends on host non-CAR CD8 T cells and tumor HLA class I integrity

As both genomic and modeling data suggest a role for eCD8T cells in mediating LBCL clearance after CD19 CAR-T infusion, we sought to functionally validate this in an immunocompetent in vivo model. Correlative studies have shown activation of non-CAR T cells in patients with clinical responses^10,11^, but whether this activation directly targets tumor cells has not been subjected to functional evaluation in a controlled setting. We therefore generated MHC-I-deficient syngeneic murine tumors through CRISPR/Cas9 deletion of *B2m* (B2*m*-KO; **Supplementary Fig. 8a,b**). An A20 clone derived from single-cell selection after sg*B2m*/CAS9 introduction lacked surface MHC-I yet retained CD19 expression and normal in-vitro proliferation (**Supplementary Fig. 8c,d**). Using m1928z^27^, we produced murine CD19-directed CAR-T cells from strain-matched Balb/C splenic T cells and co-cultured them with A20 WT and *B2m*-KO cells (**Fig. 4a**). The *B2m*-KO tumors demonstrated no impaired sensitivity to the CAR-T cells and indeed were somewhat more sensitive at lower E:T ratios, confirming that direct CAR-tumor engagement does not require target-cell MHC-I. Previous attempts to model MHC-I loss in murine LBCL models through *B2m* genomic deletion encountered problems with engraftment, hypothesized to be mediated by increased NK-cell engagement^28^. We therefore tail-vein injected WT and *B2m*-KO A20 cells to syngeneic hosts, demonstrating successful engraftment of both but with slower kinetics for the MHC-I-deficient clone (**Supplementary Fig. 8e**). Consistent with a ‘missing-self’ mechanism, purified NK cells preferentially lysed A20-B2*m*-KO targets at limiting E:T ratios while showing no difference at higher ratios (**Supplementary Fig. 8f**). Next, we assessed sensitivity of these tumors to treatment in vivo with strain-matched CD19 CAR-T cells (**Supplementary Fig. 8g**; **Fig. 4b**). WT tumors were sensitive to CAR-19 therapy, which significantly extended survival of host animals (**Fig. 4c**). *B2m*-KO tumors by contrast, despite slower engraftment and therefore a higher E:T ratio at time of CAR-T infusion, were completely insensitive to treatment (**Fig. 4d**). Thus, loss of MHC-I abrogated lymphoma clearance in an immunocompetent setting, independent of direct CAR-target recognition.

**Fig. 4.**
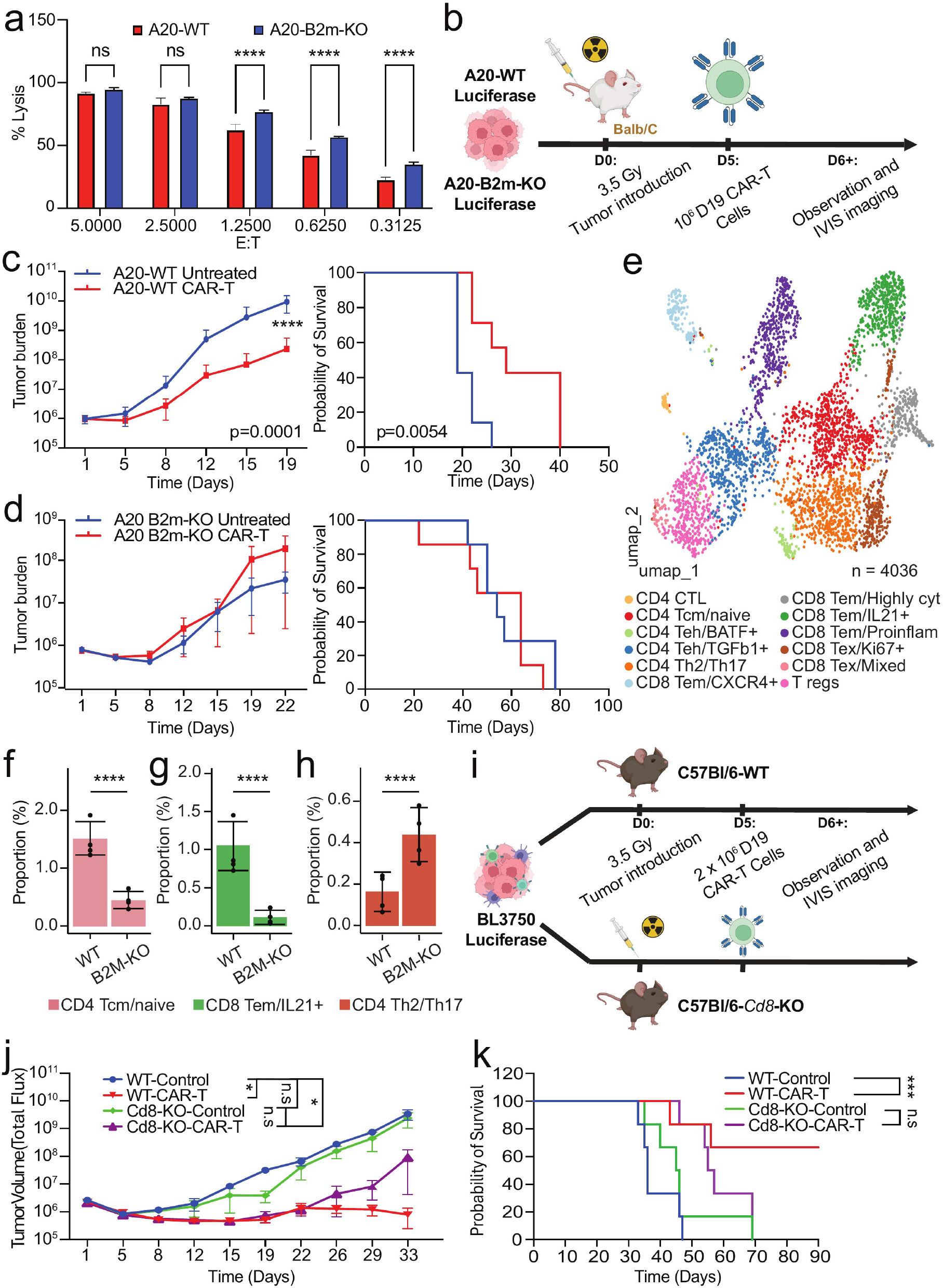
Endogenous MHC-I and CD8⁺ T cells are required for durable CAR-T control in vivo. a. Bar plot showing the tumor burden over time in mice bearing A20-WT lymphoma treated with anti-CD19 CAR T cells versus untreated controls, (**** p < 0.0001). b. Experimental design. c. Left: tumor burden (IVIS photon/sec) over time in untreated versus CAR-T-treated A20-WT mice. Right: Kaplan–Meier survival curve for untreated versus CAR-T-treated A20-WT mice (log-rank test, p = 0.0054). d. Left: tumor burden (IVIS photon/sec) over time in untreated versus CAR-T-treated A20 B2m-KO mice. Right: Kaplan–Meier survival curve for untreated versus CAR-T-treated A20 B2m-KO (log-rank test, non-significant). e. UMAP showing 4,036 T cells colored by annotated subcluster. f–h. Bar plots showing the tumor microenvironment profiling by single-cell RNA-seq, comparing A20 WT and B2m-KO tumors; p-values computed using Wilcoxon test, **** p<0.0001. i. Experimental design. j. Tumor volume over time of WT versus Cd8-KO hosts after CAR-19 treatment (n.s = non significant, * p<0.05). k. Kaplan–Meier survival curve for the four cohorts in j; p-values computed using log-rank test, *** p<0.001.

### MHC-I–deficient tumors show altered immune microenvironment and T-cell exclusion

To explore the immunologic basis of this impaired clearance, we performed single-cell RNA-seq analysis of A20 WT and *B2m*-KO tumors (n = 4 each). A total of 46,873 single cells, across eight tumors, were analyzed and clustered into malignant B cells and tumor microenvironment (TME) populations (**Supplementary Fig. 9a,b**). Both genotypes were dominated by malignant B cells (WT 86.4%, B2*m*-KO 87.2%, p = 0.023, Wilcoxon test), but WT tumors harbored more T-cell infiltration (p = 1.31e-54, Wilcoxon test), whereas B2*m*-KO tumors were enriched for monocytes (p = 1.52e-95, Wilcoxon test; **Supplementary Fig. 9c**). A total of 4,036 T cells were identified (**Fig. 4e**), distributed across subtypes (**Supplementary Fig. 9d,e**). WT tumors showed higher frequencies of immunoreactive CD4 central memory and CD8 effector memory (Tᵉᵐ) cells (p = 1.82e-28 and p = 1.38e-05 respectively, Wilcoxon test; **Fig. 4f,g**), while B2*m*-KO tumors were skewed toward immunosuppressive phenotypes, including Th2/Th17 CD4 cells (p = 3.46e-17, Wilcoxon test; **Fig. 4h**). These results reveal a relative paucity of non-CAR CD8 T-cells in the microenvironments of MHC-I-deficient tumors.

### CD8⁺ T cells are required for durable tumor clearance by CAR-19 therapy

To further evaluate the requirement for host-derived CD8⁺ T cells in CD19 CAR-T efficacy in an independent immunocompetent system, we compared CAR-T treatment of syngeneic tumors in WT and *Cd8*-KO C57BL/6 hosts (**Fig. 4i**). The MYC-driven DLBCL line BL3750^29^, expressed surface MHC-I and was sensitive to strain-matched CD19 CAR-T cells in vitro (**Supplementary Fig. 10a,b**), and engrafted with comparable kinetics in WT and *Cd8*-KO hosts in the absence of treatment (**Supplementary Fig. 10c**), indicating no inherent CD8-mediated reactivity to this tumor. Mice were then engrafted and treated with vehicle or CAR-T cells sorted to 100% CAR positivity to avoid transfer of non-CAR CD8 cells (**Supplementary Fig. 10d**). Untreated tumors engrafted in host animals with similar kinetics, leading to ethical endpoints in all animals beginning 48 days after engraftment (**Fig. 4j,k**). Tumors in *Cd8*-KO hosts, had severely compromised response to CD19 CAR-T treatment, while treated tumors in WT hosts responded well, with all mice remaining without tumor signal to the end of the experiment. Two mice in this group died during the experiment (**Fig. 4k**) but had no tumors present by IVIS imaging (**Supplementary Fig. 11**) or at necroscopy and therefore likely died related to CAR-T toxicity rather than tumor progression. Necroscopic evaluation of the remaining animals, all of whom survived until the end of the experiment at day 90, showed no evidence of tumors. All animals in the CAR-T treated CD8-KO group had tumors by IVIS imaging and grossly at necroscopy, as did all animals in both control groups. These findings confirm that clearance of LBCL tumors by CD19 CAR-T cells leading to complete elimination of tumors required the presence of eCD8 T cells in hosts to participate in therapeutic responses, validating both the genomic and mathematical model findings.

### Tumor MHC-I deficiency impairs CAR-T penetration to TMEs and secondary activation of eCD8 T cells

To functionally investigate how tumor MHC-I deficiency affects CAR-T therapy in an immunocompetent host, we injected WT or B2*m*-KO A20 cells into syngeneic mice and performed scRNA-seq on tumors with or without strain-matched m1928z CAR-T treatment, collected at day 5 to capture peak CAR-T activity (**Fig. 5a**). After quality control, 328,229 cells from 16 biologically independent tumors were retained (WT-control, n=5; WT-CAR-T, n=5; B2*m*-KO-control, n=3; B2*m*-KO-CAR-T, n=3). Unsupervised clustering resolved malignant B cells and the major immune populations of the microenvironment; the immune compartment analysis revealed substantial differences in infiltration between groups (**Fig. 5b-c** and **Supplementary Fig. 12a-c**). CAR-T therapy increased T-cell abundance in WT as expected, but this effect was absent in *B2m*-KO (p<0.05, Wilcoxon test; **Fig. 5d**). In contrast, monocyte and neutrophil populations were significantly enriched in *B2m*-KO, both with and without CAR-T treatment, suggesting inflamed myeloid-enriched TMEs known to associate with poor CAR-T outcomes in DLBCL patients (p<0.05, Wilcoxon test; **Fig. 5d**)^5,20,30^. These significant associations were also confirmed using absolute cell counts (p<0.05, Wilcoxon test; **Supplementary Fig. 13**).

**Figure 5.**
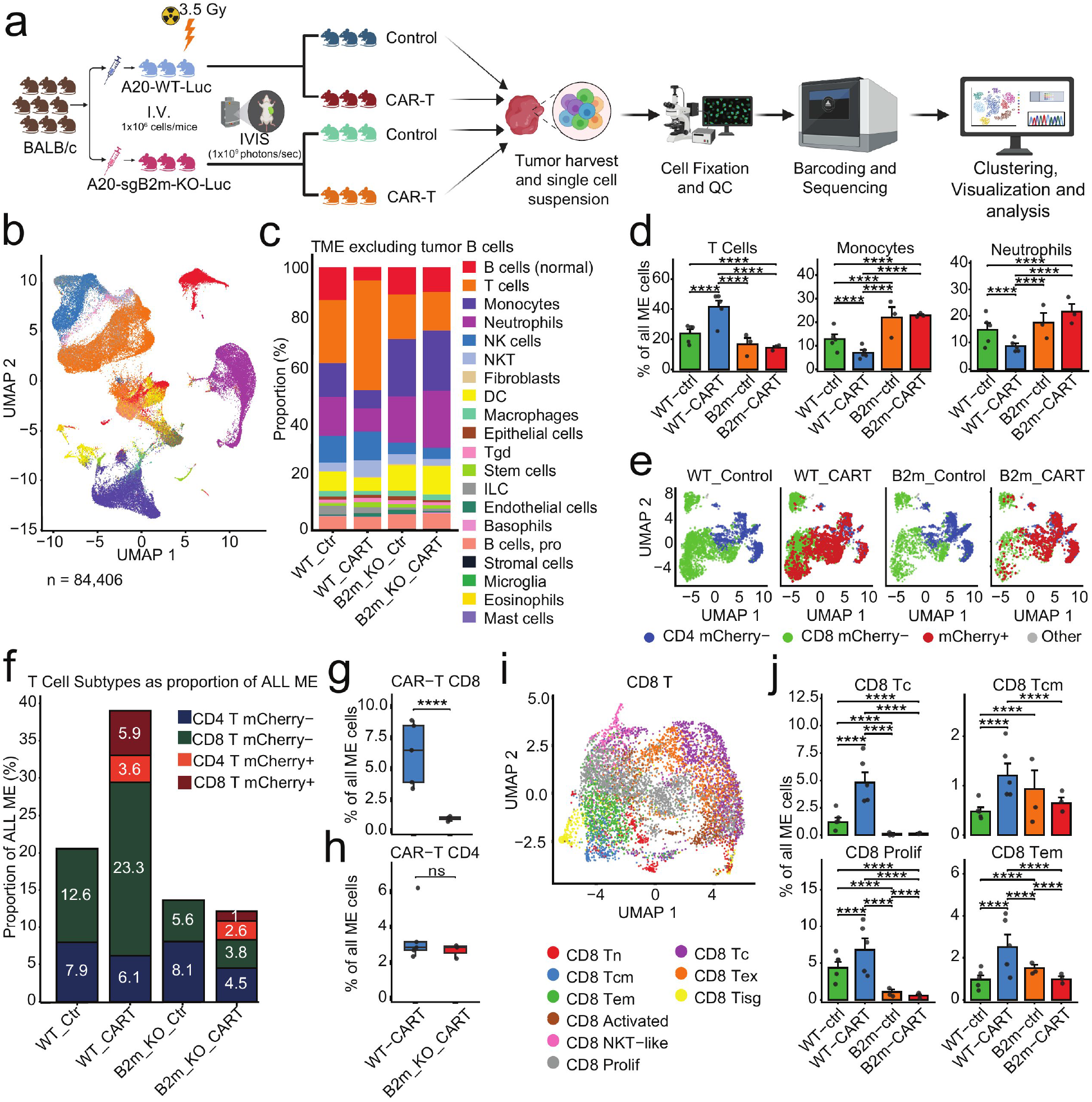
Tumor MHC-I shapes endogenous T-cell remodeling following CAR-T therapy. **a.** Experimental design for single-cell RNA sequencing of A20-WT and A20-B2m-KO tumors collected five days after anti-CD19 CAR-T therapy. **b.** UMAP of integrated single-cell transcriptomes showing major immune cell populations. **c.** Relative abundance of major cell populations across experimental groups. **d.** Proportion of T cells, monocytes and neutrophils across the immune microenvironment populations. **e.** UMAP of re-clustered T cells showing endogenous CD4, endogenous CD8, and mCherry⁺ CAR-T populations across groups. **f.** Relative abundance of endogenous and CAR-T-cell populations within the TME. **g-h.** Quantification of CD8⁺ (g) and CD4⁺ (h) CAR-T cells in WT and B2m-KO tumors. **i.** UMAP of endogenous CD8 T-cell subclusters. **j.** Quantification of cytotoxic, central-memory, proliferating, and effector-memory endogenous CD8 T-cell populations.

We therefore investigated how these TMEs impacted CAR-T penetration and activation plus secondary impacts on endogenous T-cell populations. We re-clustered 21,930 total T cells in the data and used mCherry expression to distinguish CAR from non-CAR cells in the treated samples (**Fig 5e**). These results show significant impact on especially CD8 populations mediated by the absence of MHC-I. Specifically, *B2m*-KO tumors were dramatically less infiltrated by CD8+ CAR-T cells (p<0.05, Wilcoxon test; **Fig. 5f-h**). Moreover, while eCD8 cells expanded compared to untreated in response to CAR-T in WT, they actually declined in *B2m*-KO. Examining eCD8 sub-clusters as a proportion of the non-tumor microenvironment showed deficiency in *B2m*-KO of specific populations associated with CAR-T responses, including central-memory (Tcm), effector-memory (Tem), cytotoxic (Tc), and proliferative (prolif) (p<0.05, Wilcoxon test; **Fig. 5i–j** and **Supplementary Fig. 14a-b**). Key eCD8 populations were therefore reduced in an absence of tumor MHC-I and failed to expand in response to CAR-T treatment. CAR-T therapy also altered the endogenous CD4 compartment (**Supplementary Fig. 15a-c**). Specifically, treated WT tumors contained increased proliferating and cytotoxic-like CD4 cells, whereas *B2m*-KO tumors showed relative enrichment of Th17 and regulatory T-cell populations (**Supplementary Fig. 15d**). Thus, loss of tumor MHC-I was associated with impaired endogenous CD8 expansion and a CD4 shift toward immunoregulatory states. These compositional differences were accompanied by transcriptional remodeling. In the pooled endogenous T-cell compartment, treated WT tumors expressed higher levels of cytotoxic effector genes, including *Gzmb*, *Prf1*, *Gzma*, *Serpinb6b*, and *Serpinb9*, together with *Nkg7* and *Klrk1*. By contrast, endogenous T cells from *B2m*-KO tumors preferentially expressed *Foxp3* and *Itgae,* and similar effects were seen also in the CAR-T cells themselves (**Fig. 6a**). Hallmark analysis showed prolifertive and oxidative-phosphorylation programs (E2F targets, G2M checkpoint, MYC targets, mitotic-spindle) in T cells from treated WT tumors, against IL-2-STAT5, inflammatory-response, TNF-NF-kB, TGF-beta, IL-6-JAK-STAT3, interferon-alpha, and hypoxia programs in *B2m*-KO T cells (**Fig. 6b**). Overall, tumor MHC-I associated with both the expansion and cytotoxic differentiation of endogenous T cells after CAR-T therapy. In the absence of tumor MHC-I, endogenous T cells failed to fully acquire an effector program and instead retained less differentiated and immunoregulatory features.

**Figure 6.**
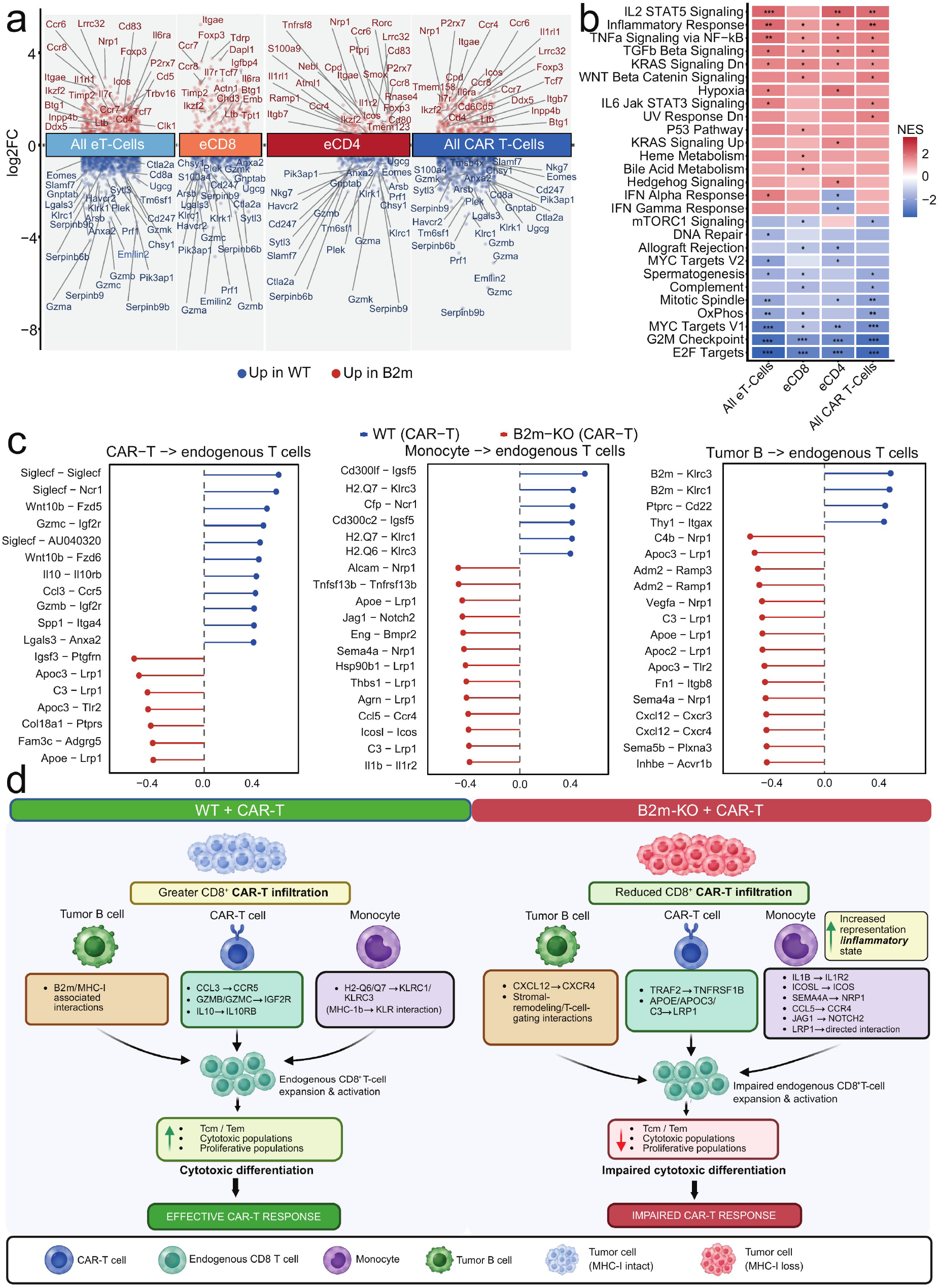
Tumor *B2m* loss remodels the CAR-T and myeloid signaling environment and impair endogenous CD8 T-cell cytotoxic differentiation. **a.** Differential gene-expression analysis comparing WT-CAR-T and B2m-KO-CAR-T pooled endogenous T cells, eCD8, eCD4, and CAR-T cells. **b.** Hallmark pathway-enrichment analysis of the corresponding T-cell populations. Positive NES indicates enrichment in B2m-KO-CAR-T and negative NES enrichment in WT-CAR-T. **c.** Differential cell–cell communication analysis using MultiNicheNet comparing CAR-T-treated WT and *B2m*-KO tumors. Differentially prioritized ligand–receptor interactions are shown for three communication axes: CAR-T cells→endogenous T cells (left), monocytes→endogenous T cells (middle), and tumor B cells→endogenous T cells (right). Blue indicates interactions preferentially prioritized in WT tumors and red indicates interactions preferentially prioritized in *B2m*-KO tumors. **d.** Integrated working model illustrating the relationship between tumor MHC-I status, CD8⁺ CAR-T infiltration, predicted intercellular communication, and endogenous CD8⁺ T-cell remodeling following CAR-T therapy.

We next examined the monocyte compartment. Sub-clustering of 13,882 monocytes resolved classical, non-classical, and intermediate subsets on the basis of canonical markers (**Supplementary Fig. 16a,b**). Differential expression between CAR-T-treated B2*m*-KO and WT tumors showed that monocytes from B2*m*-KO tumors preferentially expressed *Ccr2*, *Apoe*, *Selenop*, *Tmem176a/b*, and *Ly86*, whereas monocytes from WT tumors expressed higher *Cd38*, *Fpr1*, *Cfp*, *Ikbke*, and *Adamdec1* (**Supplementary Fig. 16c**). Consistent with this, Hallmark analysis showed enrichment of inflammatory-response, TNF-NF-kB, IL-6-JAK-STAT3, TGF-beta, and interferon-gamma programs in B2*m*-KO monocytes, with coordinate depletion of E2F targets, G2M checkpoint, MYC targets, and oxidative phosphorylation (**Supplementary Fig. 16d**). Monocytes in MHC-I-deficient tumors therefore adopt an inflammatory, less proliferative state, a phenotype we revisit below in the context of altered myeloid-to-T-cell signaling. An absence of tumor cell-surface MHC-I therefore adversely impacted CAR-T penetration to TMEs and secondary activation of endogenous T-cell populations, associated with increased myeloid infiltration and inflammatory gene expression.

MultiNicheNet^31^ prioritized differential ligand–receptor communication toward endogenous T cells from CAR-T cells, monocytes, and malignant B cells (**Fig. 6c; Supplementary Fig. 17a-g**). 1) **CAR-T→endogenous T-cells**: WT tumors prioritized Ccl3-Ccr5, granzyme-associated edges (Gzmb-Igf2r, Gzmc-Igf2r), Il10-Il10rb, among others: CCL3 is co-expressed with IFN-γ, perforin, and granzymes by effector CD8 and Th1 cells; CCR5 is recruited into the immune synapse where it costimulates CD8 priming and antitumor responses^32–34^. IL-10 is canonically anti-inflammatory but supports survival, proliferation, and cytotoxic phenotypes in antigen-activated CD8 T cells^35–36^. By contrast, B2*m*-KO tumors prioritized Traf2-Tnfrsf1b (the TNFR2-TRAF2-NF-κB module) and apolipoprotein- and complement-derived ligands converging on the scavenger receptor Lrp1 (Apoe, Apoc3, C3). TNFR2 costimulation supports Treg proliferation, stability and suppressive function, and TNFR2 upregulation on CD8 T cells precedes TOX induction and marks terminal exhaustion, while tumor-derived APOE suppresses T-cell activation^37–40^. Receiver-resolved analyses confirmed differential communication from both CD8⁺ and CD4⁺ CAR-T populations toward endogenous CD8⁺ T cells (**Supplementary Fig. 17c**). **2) Monocytes→endogenous T-cells:** WT tumors prioritized H2-Q6/H2-Q7-Klrc1/Klrc3 edges, representing MHC-Ib engagement of CD94/NKG2-family receptors. Because CD94/NKG2A recognition of non-classical MHC-Ib delivers inhibitory signals and is induced on tumor-infiltrating CD8 T cells specifically in immunologically active microenvironments^41^, this is consistent with adaptive counter-regulation in an engaged, MHC-I-visible niche. B2*m*-KO tumors instead prioritized suppressive interactions (Il1b– Il1r2, Icosl-Icos, Sema4a-Nrp1, Ccl5-Ccr4, Jag1-Notch2 and additional Lrp1-directed ligands)^42,43^, consistent with their monocytes enrichment and inflammatory transcriptional programs (TNF-NF-κB, IL6-JAK-STAT3, TGF-β and interferon programs), mirroring clinical observations that tumor interferon signaling and suppressive myeloid populations associate with CAR-T failure in large B-cell lymphoma^5^(**Supplementary Fig. 17d**). **3) Tumor-B-cell→endogenous T-cells**: Dramatic prioritization of B2*m* edges in WT is an obligate consequence of comparison to B2*m*-KO, providing internal validation for the analysis. B2*m*-KO by contrast prioritized stromal-remodeling and T-cell-gating interactions, including CXCL12-CXCR4, a canonical mediator of T-cell exclusion whose blockade increases intratumoral T-cell infiltration^44,45^. These predicted interactions therefore provide a mechanistic correlate for the reduced CD8⁺ CAR-T infiltration and the failure of endogenous CD8⁺ expansion that we measured in B2*m*-KO tumors (**Fig. 6d**; **Supplementary Fig. 17e**).

### CAR-T cell therapy enhances anti-tumoral response of endogenous T cells

These data implicate intact MHC-I and host eCD8⁺ T-cell participation in CD19 CAR-T efficacy, prompting us to test whether CAR-T therapy promotes acquisition of endogenous antitumor reactivity. We therefore performed interferon-gamma (IFNγ) ELISPOT assays using patient-derived effector cells collected before and after CAR-T infusion, testing reactivity against autologous tumor targets as previously described by Neelapu et al^1,46^. Baseline samples consisted of cryopreserved peripheral blood mononuclear cells (PBMCs), leukapheresis products, or tumor-infiltrating lymphocytes (TILs) collected prior to CAR-T infusion, depending on availability. Post-infusion samples consisted of cryopreserved PBMCs collected between 40 and 350 days after CAR-T (**Supplementary Table 8**). To specifically assess the activity of the endogenous T cell compartment post-infusion, CAR+ T cells were depleted from these post-treatment PBMC samples prior to assessment. All samples were co-cultured with autologous tumor cell suspensions that had been pre-depleted of T cells and activated with CD40L and IL-4 (**Fig. 7a**). Post-infusion samples showed significantly increased IFNγ production in response to autologous tumor challenge compared to baseline, indicating enhanced tumor-specific reactivity of the endogenous T cell compartment consistent with CAR-T– mediated epitope spreading. (p=0.039, two-way ANOVA; **Fig. 7b**; **Supplementary Fig. 18)**.

**Fig. 7.**
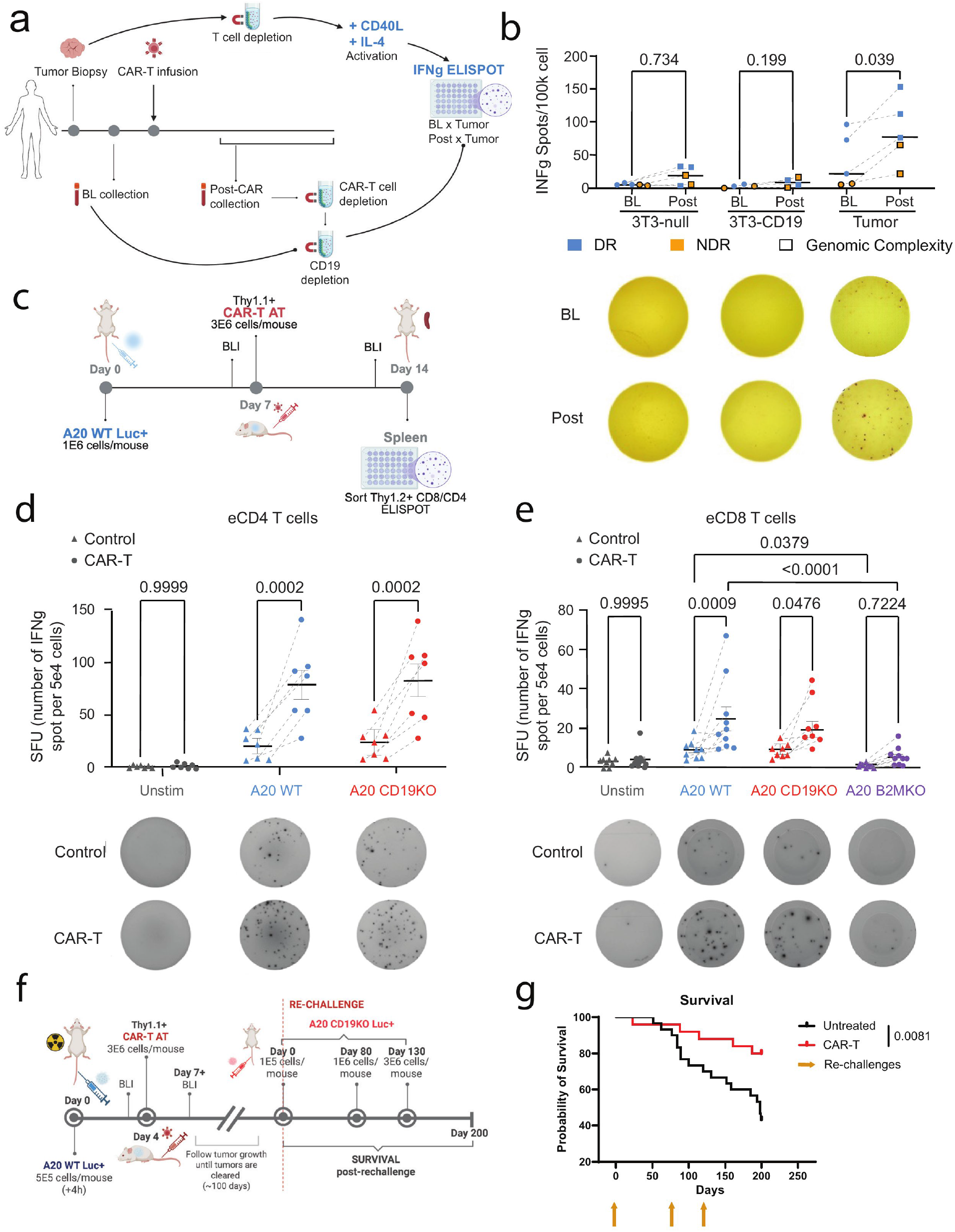
CAR-T cell therapy enhances anti-tumoral response of endogenous T cells. **a.** Schematic representation of the workflow followed to collect and process LBCL patient samples to quantify anti-tumor response measured by IFNγ ELISPOT. Briefly, surgical excisional tumor biopsies were processed to generate viable cell suspensions, depleted from T cells and activated with CD40L and IL4. Pre-(Baseline, BL) and post-CAR-T infusion samples were collected, PBMCs enriched, and CAR+ T cells depleted from Post-CAR-T samples prior co-culture. BL and post samples were co-cultured with activated tumors and their respective controls on ELISPOT plates for 48h before revealing spots. **b.** IFNγ spot forming units (SFU) per 100,000 cells by ELISPOT from pre-(BL) and post-CAR-T-infusion patient samples (n=5) co-cultured with autologous tumor biopsies cell suspensions, or 3T3-null and 3T3-hCD19+ as controls. **c.** Schematic representation of murine validation experimental design. A20 tumor-bearing Balb\c mice were treated with Thy1.1+ CAR-T cells. Two weeks post-tumor challenge, splenic endogenous (Thy1.2^+^ Thy1.1^-^ CAR^-^) CD4 or CD8 T cells were FACS sorted and co-cultured for 24h with different A20 target cells (E:T 1:1) on IFNγ ELISPOT plates. **d-e.** Number of IFNγ SFU per 50,000 cells of eCD4 (**d**) or eCD8 (**e**) unstimulated (unstim), or stimulated with A20 WT, CD19KO or B2MKO (MHC-I KO, eCD8 only). Data was pooled from three independent experiments (n=10). Statistical analysis was performed using a Two-way ANOVA. **f.** schematic representation of rechallenge experiment. Briefly, BALB/c mice were inoculated with A20 WT CD19⁺ luciferase-expressing lymphoma cells (5×10⁵ cells/mouse) and treated with 3×10⁶ CD19-directed CAR-T cells following confirmation of tumor engraftment and normalization of tumor burden. Mice achieving durable tumor clearance were rechallenged 100 days after the initial tumor inoculation with A20 CD19KO luciferase-expressing lymphoma cells lacking expression of the CAR target antigen. In three independent experiments, surviving mice underwent sequential CD19KO tumor rechallenges at escalating doses (1×10⁵, 1×10⁶, and 3×10⁶ cells/mouse). Age-matched naïve BALB/c mice receiving identical tumor challenges served as untreated controls. **g)** Kaplan-Meier survival analysis of pooled data from all experiments demonstrated significantly improved survival among previously CAR-T-treated mice compared with untreated controls (log-rank p = 0.0081). Protection against repeated challenge with CD19-deficient tumors suggests that CAR-T-mediated tumor elimination generated durable endogenous antitumor immune responses directed against antigens beyond the CAR target and capable of mediating long-term tumor control; orange arrows indicate the days on which re-challenge was performed.

We next further investigated this phenomenon in a controlled preclinical setting (**Fig. 7c**). We injected Thy1.2+ Balb/C mice intravenously (i.v.) with luciferase-expressing A20 WT cells. One week later, we stratified tumor burden by bioluminescent signal randomized the animals to receive i.v. injections of Thy1.1^+^ CD19-targeted CAR-T cells (CAR-T group) or no treatment (tumor-only controls) and harvested spleens seven days later. Endogenous Thy1.2^+^ CD4^+^ (eCD4) and eCD8 T cells were fluorescence-activated cell sorting (FACS)–purified (**Supplementary Fig. 19a**) and co-cultured with different A20 tumor cells: (i) A20 WT; (ii) A20 *Cd19*-knockout (*Cd19*-KO) to exclude residual CAR-T-mediated activation; and (iii) MHC class I–deficient tumor cells (*B2m*-KO) to abrogate T cell receptor (TCR)-mediated activation (**Supplementary Fig. 19b**). After 24h, IFNγ ELISPOT plates were read and spot-forming units (SFU) quantified compared to unstimulated (Unstim.) controls, to which no tumor cells were added. Consistent with observations in LBCL patients, both eCD4 and eCD8 T cells from CAR-T–treated mice exhibited a significant increase in IFNγ production compared to tumor-only controls (p<0.05, two-way ANOVA; **Fig. 7d,e**; **Supplementary Fig. 19c,d**; **Supplementary Table 9**). This enhanced response persisted when T cells were challenged with *Cd19*-KO tumor cells. Furthermore, eCD8 T cells co-cultured with *B2m*-KO A20 cells showed no significant increase in IFNγ spots relative to those challenged with MHC-I–expressing tumors (p=0.72, two-way ANOVA; **Fig. 7e**), implicating TCR-MHC-I immune synapse formation as a requirement for acquired tumor reactivity. Together, these findings suggest that CD19-directed CAR-T therapy can facilitate the expansion and activation of endogenous T cells against tumors independent of the CD19 CAR target. To determine whether this acquired endogenous antitumor reactivity translated into durable protection against antigen-loss tumor variants in vivo, we performed sequential rechallenge experiments using A20 lymphoma cells lacking the CAR target antigen CD19. Balb/c mice bearing luciferase-expressing A20 WT tumors were treated with CD19-directed CAR-T cells and monitored until complete tumor clearance. Mice with durable remissions were rechallenged with A20 CD19KO lymphoma cells 100 days after the initial tumor inoculation. Across three independent experiments, surviving mice underwent a total of three sequential CD19KO tumor challenges at escalating doses (**Fig. 7f**). Age-matched naïve Balb/c mice receiving identical tumor inoculations served as untreated controls. Despite the absence of the CAR target antigen, previously CAR-T-treated mice exhibited significantly prolonged survival compared with control animals following repeated CD19KO tumor challenge (p=0.0081, log-rank test; **Fig. 7g**). The ability of CAR-T-cured mice to control tumors lacking CD19 indicates the development of durable host-derived antitumor immunity directed against antigens beyond the CAR target. Together with the ELISPOT findings, these results demonstrate that CD19 CAR-T therapy induces durable endogenous antitumor immunity capable of mediating protection against antigen-loss tumor variants, supporting a model in which CAR-T treatment broadens host antitumor immune recognition beyond CD19.

## DISCUSSION

Adoptive transfer of CD19-directed CAR T cells has transformed outcomes for relapsed/refractory LBCL, yet a substantial proportion of patients relapse or fail to respond despite preserved CD19 expression^1,2,14,15^. Here, by integrating whole-genome and transcriptome profiling with functional patient assays, syngeneic models, and mathematical modeling, the data demonstrate how durable CAR-19 responses require not only target expression but also intact tumor immunogenicity and engagement of endogenous, non-CAR CD8⁺ T cells. Together these results position CAR-19 as both a tumor-directed therapy and a catalyst for durable host antitumor immunity that extends beyond the CAR target itself. Tumor-cell escape from eCD8 cells is therefore a novel resistance driver associated with poor outcome. This model is supported by multiple lines of evidence. First, we showed that disruption of immune-synapse components usually in a context of genomic complexity associate with early progression after CART, independently from tumor burden and CD19 expression. Second, a dynamical math model of CAR-T, previously parameterized on different patient data, supported the biological plausibility of cooperation between CAR-T (engaging CD19) and eCD8 (engaging MHC-I) to effect response, recapitulating our patient data: without loss of endogenous immunogenicity, higher pre-treatment NeoAg associates with response. Third, MHC-I-deficient (*B2m*-KO) tumors remained fully sensitive to CAR-T killing in vitro yet were insensitive in vivo in immunocompetent hosts, despite slower engraftment kinetics, indicating that direct CAR-target binding is insufficient for clearance when tumors evade host T-cell recognition. Fourth, an absence of CD8⁺ T cells prevented CAR-T sustained efficacy against an immunocompetent LBCL model, while wild-type hosts achieved durable remissions with no residual tumor at necropsy. Complementing these models, patient and murine ELISPOT assays showed enhanced post-infusion reactivity of endogenous T cells against autologous tumor, including against CD19-KO targets. Consistent with this, mice cured by CD19 CAR-T therapy controlled sequential rechallenge with CD19-deficient A20 tumors at escalating doses, surviving significantly longer than age-matched naïve controls, demonstrating that the host-derived immunity generated during CAR-T-mediated clearance is durable and functionally protective against antigen-loss variants. These results are consistent with a model where CART potentially facilitate both pre-existing host immunogenicity against the tumor and potential epitope spreading, although we cannot implicate specific tumor neoantigens without further extensive investigation. Single-cell RNA-seq further supported this interpretation by revealing greater T-cell infiltration and immunoreactive CD8 T-cell states in MHC-I-competent tumors relative to MHC-I-deficient tumors. Importantly, ELISPOT assays performed on samples from the same patients with available WGS and RNA-seq data suggested concordance between high immunogenicity, intact HLA expression, and underlying genomic complexity.

Overall, these findings refine, and to some extent reframe, the predominant resistance narrative in LBCL. While high disease burden promoting immune exhaustion and reduction of CD19 undoubtedly explains a subset of failures^47^, our results indicate that CAR target-independent mechanisms centered on escape from the host immune surveillance play a key role. Patient genomics, controlled in vivo functional experiments, and computational modeling collectively demonstrated that effective tumor eradication requires CAR T cells and the host immune system to act in concert. Therefore, novel treatment strategies should aim not only to maximize CAR T-mediated cytotoxicity but also to enhance their ability to stimulate host immune responses.

## METHODS

Germline and tumor samples were collected prospectively under institutional review board protocols^6^ and in accordance with the Declaration of Helsinki. Details of the analysis populations, whole-genome and transcriptome sequencing, computational modeling, single-cell RNA sequencing, mouse models and statistical analysis are provided in **Supplementary Methods**.

## DATA AVAILABILITY

The WGS and RNA-seq from biopsies of CAR-19–treated patients have been deposited at the Database of Genotypes and Phenotypes with the study accession phs003023.v2.p1.

## AUTHOR CONTRIBUTIONS

F.M., J.H.S., A.R.A.A., F.L.L., designed and supervised the study, collected, generated and analyzed the data and wrote the paper. B.Z., C.S.F., V.V.G.S., C.G., generated, analyze and interpreted the data and wrote the paper. A.A.Y., M.C., M.M., L.C.D., M.D., K.R., J.A.M., S.C., E.M., W.S., A.A., A.D.N., D.C., C.A., J.Y.S., M.D., O.L., J.P.A., M.R.T., R.A., S.U., generated and analyze the data. J.M., M.D.J., collected the samples and clinical data

## ACKNOWLEDGMENTS

This work was supported by the Sylvester Comprehensive Cancer Center National Cancer Institute Core Grant (P30 CA 240139), Memorial Sloan Kettering Cancer Center National Cancer Institute Core Grant (P30 CA 008748) and Moffitt Cancer Center National Cancer Institute Core Grant (P30CA076292)

F.L.L. is supported in part by the Leukemia and Lymphoma Society as a Clinical Scholar, the National Cancer Institute (R01CA244328-03), and generous donations from the Hano, Hyer family, Hoenle Family and the Thiel family.

FM is supported by NIH-NCI (R37CA289752), Myeloma Solution Funds, Comedy vs Cancer, Cycle for Survival, and by the Paula and Rodger Riney Foundation.

This work was supported by the US Department of Defense Congressionally Directed Medical Research Program (CDMRP) under award GR023242 to J.H.S.

## REFERENCES

1 Neelapu, S. S. et al. Axicabtagene Ciloleucel CAR T-Cell Therapy in Refractory Large B-Cell Lymphoma. N Engl J Med 377, 2531–2544 (2017). 10.1056/NEJMoa1707447

2 Abramson, J. S. et al. Lisocabtagene maraleucel for patients with relapsed or refractory large B-cell lymphomas (TRANSCEND NHL 001): a multicentre seamless design study. Lancet 396, 839–852 (2020). 10.1016/S0140-6736(20)31366-0

3 Cherng, H. J. et al. Risk assessment with low-pass whole-genome sequencing of cell-free DNA before CD19 CAR T-cell therapy for large B-cell lymphoma. Blood 140, 504–515 (2022). 10.1182/blood.2022015601

4 Deng, Q. et al. Characteristics of anti-CD19 CAR T cell infusion products associated with efficacy and toxicity in patients with large B cell lymphomas. Nat Med 26, 1878–1887 (2020). 10.1038/s41591-020-1061-7

5 Jain, M. D. et al. Tumor interferon signaling and suppressive myeloid cells are associated with CAR T-cell failure in large B-cell lymphoma. Blood 137, 2621–2633 (2021). 10.1182/blood.2020007445

6 Jain, M. D. et al. Whole-genome sequencing reveals complex genomic features underlying anti-CD19 CAR T-cell treatment failures in lymphoma. Blood 140, 491–503 (2022). 10.1182/blood.2021015008

7 Sworder, B. J. et al. Determinants of resistance to engineered T cell therapies targeting CD19 in large B cell lymphomas. Cancer Cell 41, 210–225 e215 (2023). 10.1016/j.ccell.2022.12.005

8 Shouval, R. et al. Cytokine Release Syndrome and Neurotoxicity Following CD19 CAR-T in B-Cell Lymphoma. Transplant Cell Ther 31, 419–433 (2025). 10.1016/j.jtct.2025.03.011

9 Yu, H. et al. Repeated loss of target surface antigen after immunotherapy in primary mediastinal large B cell lymphoma. Am J Hematol 92, E11–E13 (2017). 10.1002/ajh.24594

10 Cheloni, G. et al. Durable response to CAR T is associated with elevated activation and clonotypic expansion of the cytotoxic native T cell repertoire. Nat Commun 16, 4819 (2025). 10.1038/s41467-025-59904-x

11 Chen, P. H. et al. Activation of CAR and non-CAR T cells within the tumor microenvironment following CAR T cell therapy. JCI Insight 5 (2020). 10.1172/jci.insight.134612

12 Alizadeh, D. et al. IFNgamma Is Critical for CAR T Cell-Mediated Myeloid Activation and Induction of Endogenous Immunity. Cancer Discov 11, 2248–2265 (2021). 10.1158/2159-8290.CD-20-1661

13 Sanmamed, M. F. et al. A Burned-Out CD8(+) T-cell Subset Expands in the Tumor Microenvironment and Curbs Cancer Immunotherapy. Cancer Discov 11, 1700–1715 (2021). 10.1158/2159-8290.CD-20-0962

14 Locke, F. L. et al. Long-term safety and activity of axicabtagene ciloleucel in refractory large B-cell lymphoma (ZUMA-1): a single-arm, multicentre, phase 1-2 trial. Lancet Oncol 20, 31–42 (2019). 10.1016/S1470-2045(18)30864-7

15 Schuster, S. J. et al. Tisagenlecleucel in Adult Relapsed or Refractory Diffuse Large B-Cell Lymphoma. N Engl J Med 380, 45–56 (2019). 10.1056/NEJMoa1804980

16 Alexandrov, L. B. et al. The repertoire of mutational signatures in human cancer. Nature 578, 94–101 (2020). 10.1038/s41586-020-1943-3

17 Nik-Zainal, S. et al. Landscape of somatic mutations in 560 breast cancer whole-genome sequences. Nature 534, 47–54 (2016). 10.1038/nature17676

18 Zhang, Z. et al. Point mutation in CD19 facilitates immune escape of B cell lymphoma from CAR-T cell therapy. J Immunother Cancer 8 (2020). 10.1136/jitc-2020-001150

19 Plaks, V. et al. CD19 target evasion as a mechanism of relapse in large B-cell lymphoma treated with axicabtagene ciloleucel. Blood 138, 1081–1085 (2021). 10.1182/blood.2021010930

20 Locke, F. L. et al. Impact of tumor microenvironment on efficacy of anti-CD19 CAR T cell therapy or chemotherapy and transplant in large B cell lymphoma. Nat Med 30, 507–518 (2024). 10.1038/s41591-023-02754-1

21 Spiegel, J. Y. et al. CAR T cells with dual targeting of CD19 and CD22 in adult patients with recurrent or refractory B cell malignancies: a phase 1 trial. Nat Med 27, 1419–1431 (2021). 10.1038/s41591-021-01436-0

22 Mardis, E. R. Neoantigens and genome instability: impact on immunogenomic phenotypes and immunotherapy response. Genome Med 11, 71 (2019). 10.1186/s13073-019-0684-0

23 Mandal, R. et al. Genetic diversity of tumors with mismatch repair deficiency influences anti-PD-1 immunotherapy response. Science 364, 485–491 (2019). 10.1126/science.aau0447

24 MacArthur, R. Species packing and competitive equilibrium for many species. Theor Popul Biol 1, 1–11 (1970). 10.1016/0040-5809(70)90039-0

25 Berec, L. Impacts of foraging facilitation among predators on predator-prey dynamics. Bull Math Biol 72, 94–121 (2010). 10.1007/s11538-009-9439-1

26 Kimmel, G. J., Locke, F. L. & Altrock, P. M. The roles of T cell competition and stochastic extinction events in chimeric antigen receptor T cell therapy. Proc Biol Sci 288, 20210229 (2021). 10.1098/rspb.2021.0229

27 Davila, M. L., Kloss, C. C., Gunset, G. & Sadelain, M. CD19 CAR-targeted T cells induce long-term remission and B Cell Aplasia in an immunocompetent mouse model of B cell acute lymphoblastic leukemia. PLoS One 8, e61338 (2013). 10.1371/journal.pone.0061338

28 Fangazio, M. et al. Genetic mechanisms of HLA-I loss and immune escape in diffuse large B cell lymphoma. Proc Natl Acad Sci U S A 118 (2021). 10.1073/pnas.2104504118

29 Minard-Colin, V. et al. Lymphoma depletion during CD20 immunotherapy in mice is mediated by macrophage FcgammaRI, FcgammaRIII, and FcgammaRIV. Blood 112, 1205–1213 (2008). 10.1182/blood-2008-01-135160

30 Scholler, N. et al. Tumor immune contexture is a determinant of anti-CD19 CAR T cell efficacy in large B cell lymphoma. Nat Med 28, 1872–1882 (2022). 10.1038/s41591-022-01916-x

31 Browaeys, R. et al. (2023). 10.1101/2023.06.13.544751

32 Gonzalez-Martin, A., Mira, E. & Manes, S. CCR5 in cancer immunotherapy: More than an “attractive” receptor for T cells. Oncoimmunology 1, 106–108 (2012). 10.4161/onci.1.1.17995

33 Kang, T. G., Park, H. J., Moon, J., Lee, J. H. & Ha, S. J. Enriching CCL3 in the Tumor Microenvironment Facilitates T cell Responses and Improves the Efficacy of Anti-PD-1 Therapy. Immune Netw 21, e23 (2021). 10.4110/in.2021.21.e23

34 Nicolet, B. P. et al. CD29 identifies IFN-gamma-producing human CD8(+) T cells with an increased cytotoxic potential. Proc Natl Acad Sci U S A 117, 6686–6696 (2020). 10.1073/pnas.1913940117

35 Xi, J. et al. Stimulatory role of interleukin 10 in CD8(+) T cells through STATs in gastric cancer. Tumour Biol 39, 1010428317706209 (2017). 10.1177/1010428317706209

36 Naing, A. et al. PEGylated IL-10 (Pegilodecakin) Induces Systemic Immune Activation, CD8(+) T Cell Invigoration and Polyclonal T Cell Expansion in Cancer Patients. Cancer Cell 34, 775–791 e773 (2018). 10.1016/j.ccell.2018.10.007

37 Hoyt-Miggelbrink, A. M. et al. Upregulation of TNFR2 Precedes TOX Expression by Exhausted T cells and Restricts Antitumor and Antiviral Immunity. Clin Cancer Res 32, 782–800 (2026). 10.1158/1078-0432.CCR-24-3455

38 Lubrano di Ricco, M., et al. Tumor necrosis factor receptor family costimulation increases regulatory T-cell activation and function via NF-kappaB. Eur J Immunol 50, 972–985 (2020). 10.1002/eji.201948393

39 Sizova, O., John, L. S., Ma, Q. & Molldrem, J. J. Multi-faceted role of LRP1 in the immune system. Front Immunol 14, 1166189 (2023). 10.3389/fimmu.2023.1166189

40 Wu, X. et al. Tumor Apolipoprotein E is a key checkpoint blocking anti-tumor immunity in mouse melanoma. Front Immunol 13, 991790 (2022). 10.3389/fimmu.2022.991790

41 Lohwasser, S., Kubota, A., Salcedo, M., Lian, R. H. & Takei, F. The non-classical MHC class I molecule Qa-1(b) inhibits classical MHC class I-restricted cytotoxicity of cytotoxic T lymphocytes. Int Immunol 13, 321–327 (2001). 10.1093/intimm/13.3.321

42 Chen, L. et al. IL1R2 increases regulatory T cell population in the tumor microenvironment by enhancing MHC-II expression on cancer-associated fibroblasts. Journal for ImmunoTherapy of Cancer 10 (2022). 10.1136/jitc-2022-004585

43 Martin-Orozco, N. et al. Melanoma cells express ICOS ligand to promote the activation and expansion of T-regulatory cells. Cancer Res 70, 9581–9590 (2010). 10.1158/0008-5472.CAN-10-1379

44 Deipenbrock, A. et al. Targeting stroma-mediated T-cell exclusion and functional exhaustion in pancreatic ductal adenocarcinoma through CXCR4 and PD-1 blockade. Front Immunol 17, 1844781 (2026). 10.3389/fimmu.2026.1844781

45 Zboralski, D., Hoehlig, K., Eulberg, D., Fromming, A. & Vater, A. Increasing Tumor-Infiltrating T Cells through Inhibition of CXCL12 with NOX-A12 Synergizes with PD-1 Blockade. Cancer Immunol Res 5, 950–956 (2017). 10.1158/2326-6066.CIR-16-0303

46 Malyguine, A. et al. A modified human ELISPOT assay to detect specific responses to primary tumor cell targets. J Transl Med 2, 9 (2004). 10.1186/1479-5876-2-9

47 Ruella, M., Korell, F., Porazzi, P. & Maus, M. V. Mechanisms of resistance to chimeric antigen receptor-T cells in haematological malignancies. Nat Rev Drug Discov 22, 976–995 (2023). 10.1038/s41573-023-00807-1

