## Supplementary Figures for "Lymphoma response to chimeric antigen receptor T cell therapy is driven by endogenous tumor-reactive CD8⁺ T cells"

### SUPPLEMENTARY FIGURE LEGENDS

**Supplementary Fig. 1. Analysis populations**. Flow of patients from the sequenced cohort to each analysis population. Boxes give the number of patients contributing to each analysis; annotations on the right give the number excluded at each step and the reason.

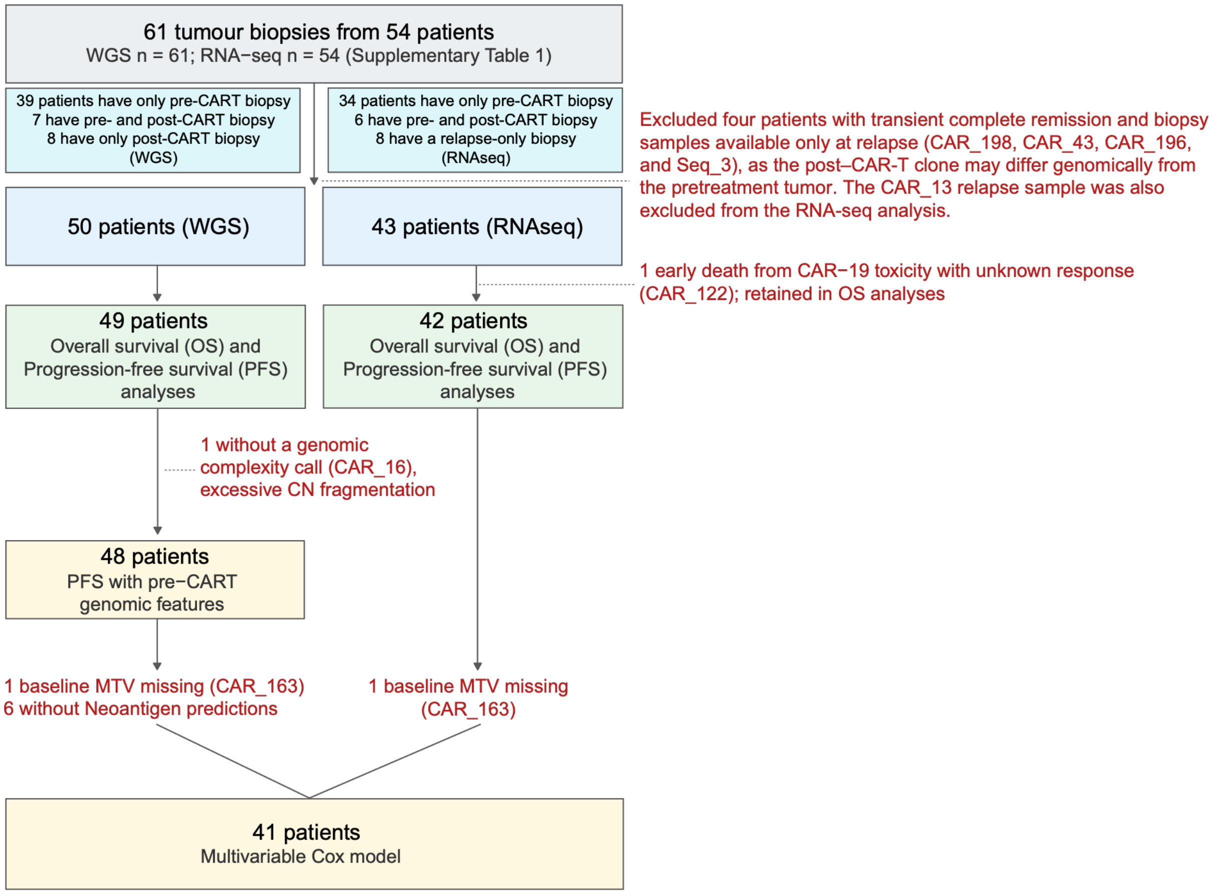

**Supplementary Fig. 2. Genomic features associated with poor outcome after CD19 CAR-T treatment. a-f.** PFS Kaplan-Meier curves per each significant genomic driver (p value obtained using log-Rank test). **g.** Cox multivariate model between the significant genomic drivers and important clinical features.

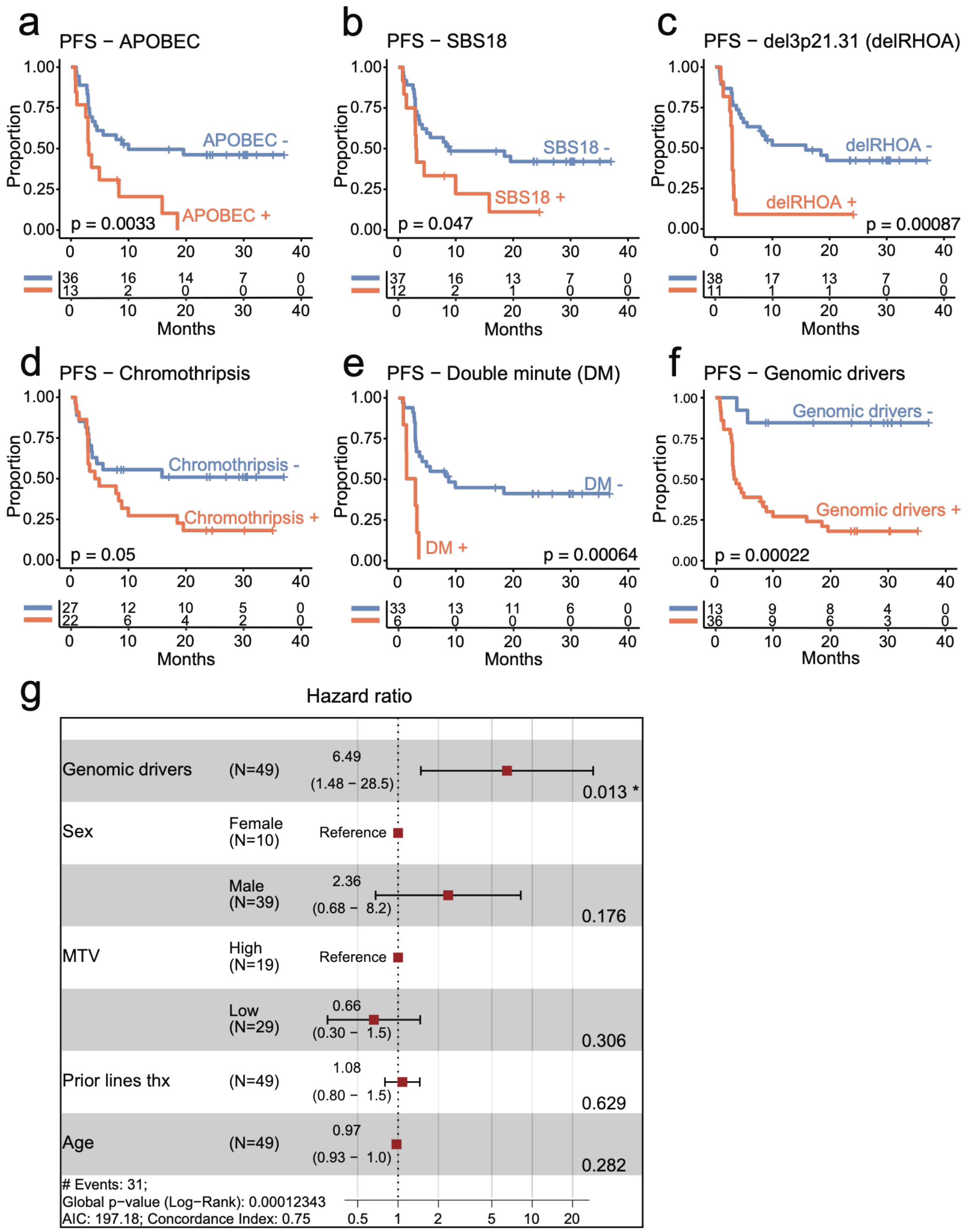

**Supplementary Fig. 3. Complex signature analysis. a.** Analysis workflow**. b-c.** Extracted CNV (**b**) and SV (**c**) signatures using HDP, each bar plot shows the contribution of each feature into each signature.

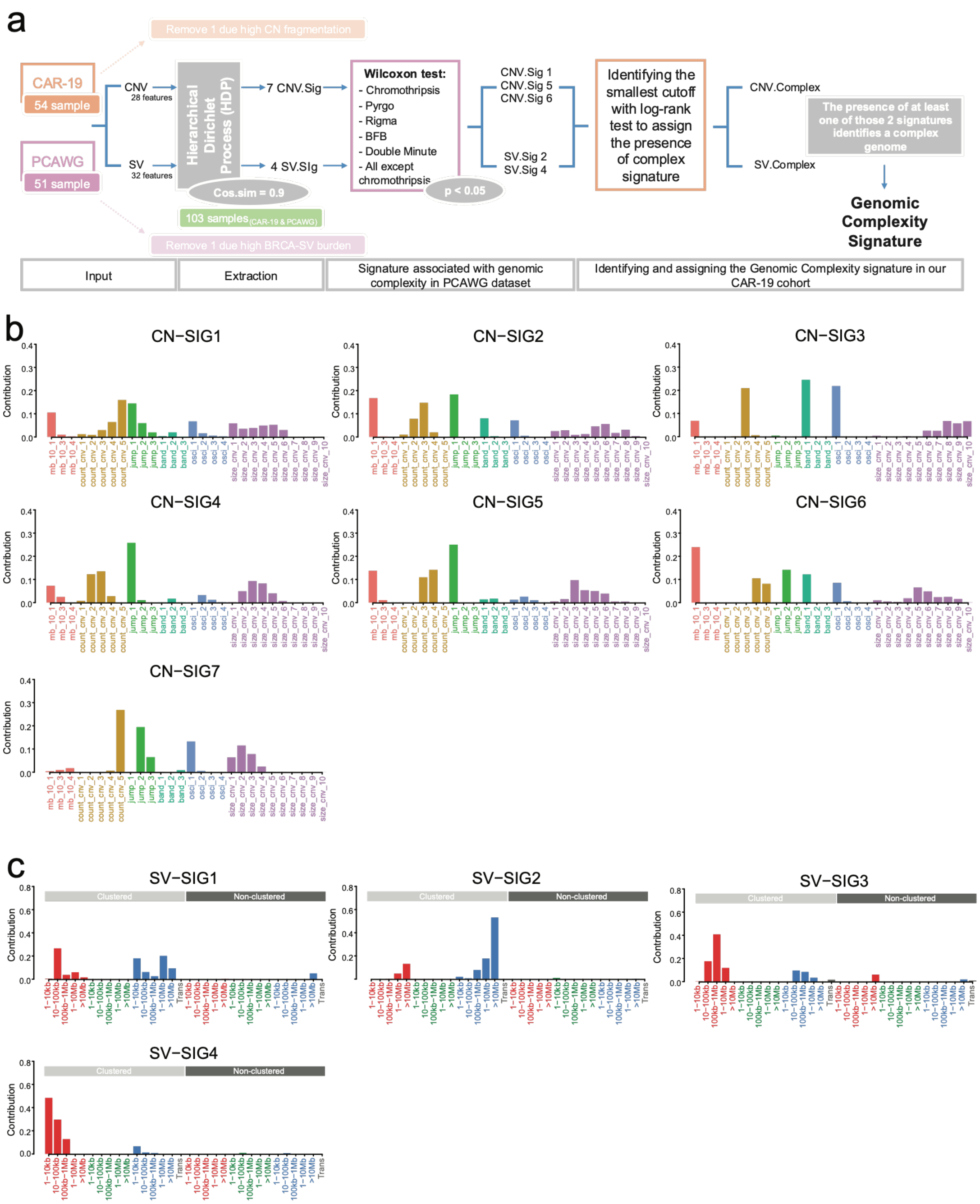

**Supplementary Fig. 4. Phylogenetic analysis. a-g.** Phylogenetic trees. Y-axes are the number of variants. The violet line represents the trunk, while the red, blue, yellow and green are the branches, when present. The cartoons indicate the biopsy location

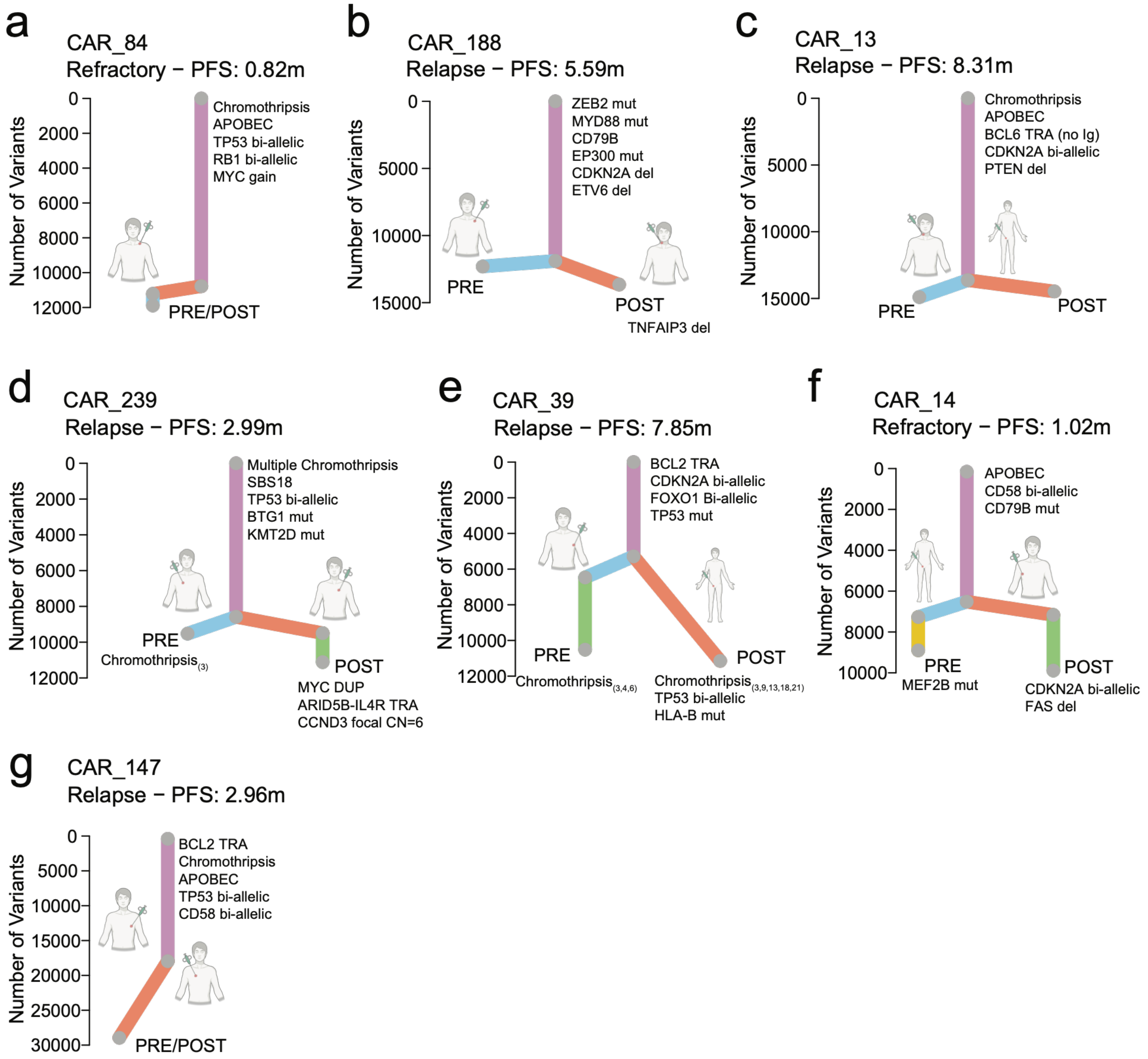

**Supplementary Fig. 5. Neoantigen burden and Immune synapse genomic analysis workflow.** In yellow somatic variant analysis; in blue MHC class I calls; in green neoantigen burden flow.

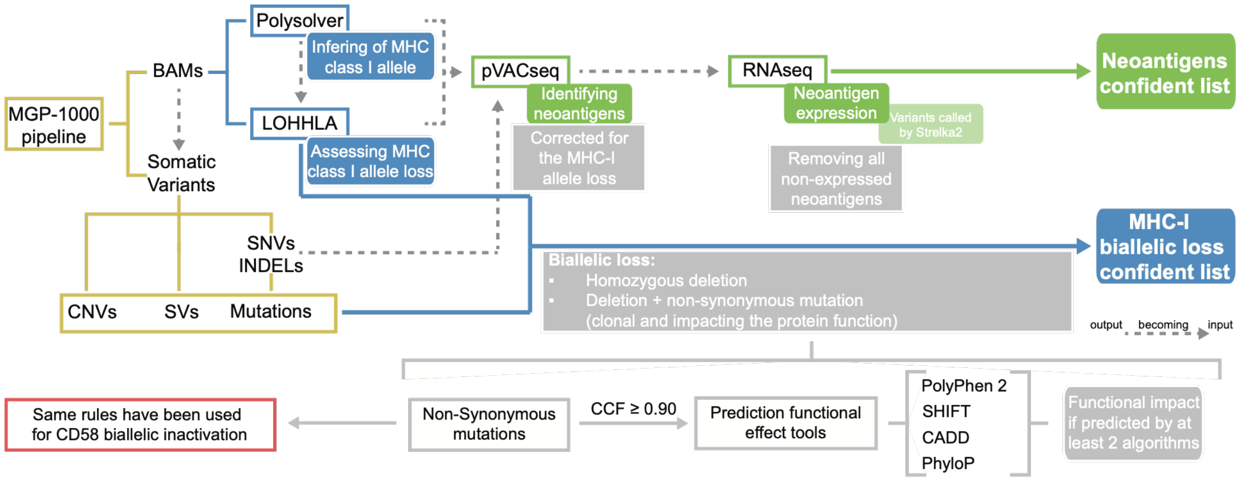

**Supplementary Fig. 6. Neoantigen burden. a.** Predicted clonal neoantigen burden by clinical outcome (Durable Responders (DR), blue; Non-Durable Responders (NDR), orange); no significant difference in the unstratified cohort (two-sided Wilcoxon, p = 0.7615). Each dot represents a patient, patients harboring immune-evasion features are shown in red, wild-type in grey. **b.** Predicted clonal neoantigen burden by genomic complexity; no significant difference in the unstratified cohort (two-sided Wilcoxon). **c.** Bar plot showing enrichment of neoantigen burden in patients with biallelic inactivation of immune-synapse (IS) genes (HLA-A, HLA-B, HLA-C, B2M and CD58); in blue patients harboring IS biallelic inactivation, in grey patients with IS WT; p-value computed using Fisher's Exact test (WT: wild type).

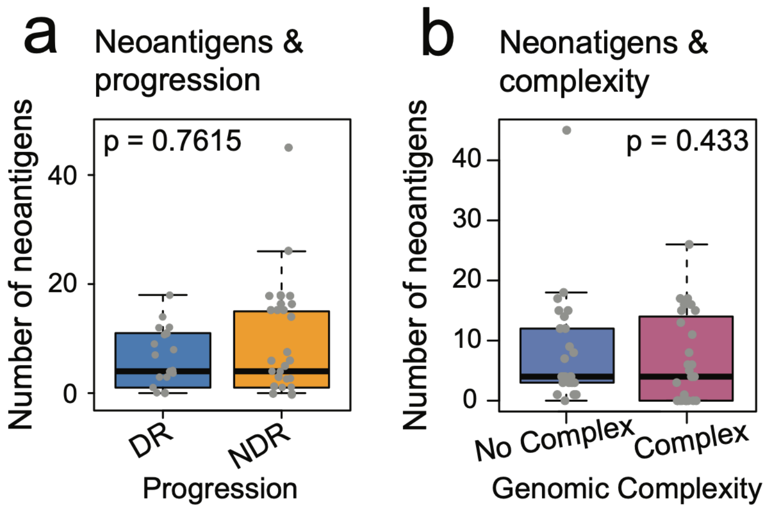

**Supplementary Fig. 7. Immune-evasion mechanism events. a-n**. CNVs/SVs profiles for MHC-I complex and CD58. Vertical lines: Red = deletion, blue = inversion, green = duplications, black = translocation. Gene names: violet = mutated, red = WT.

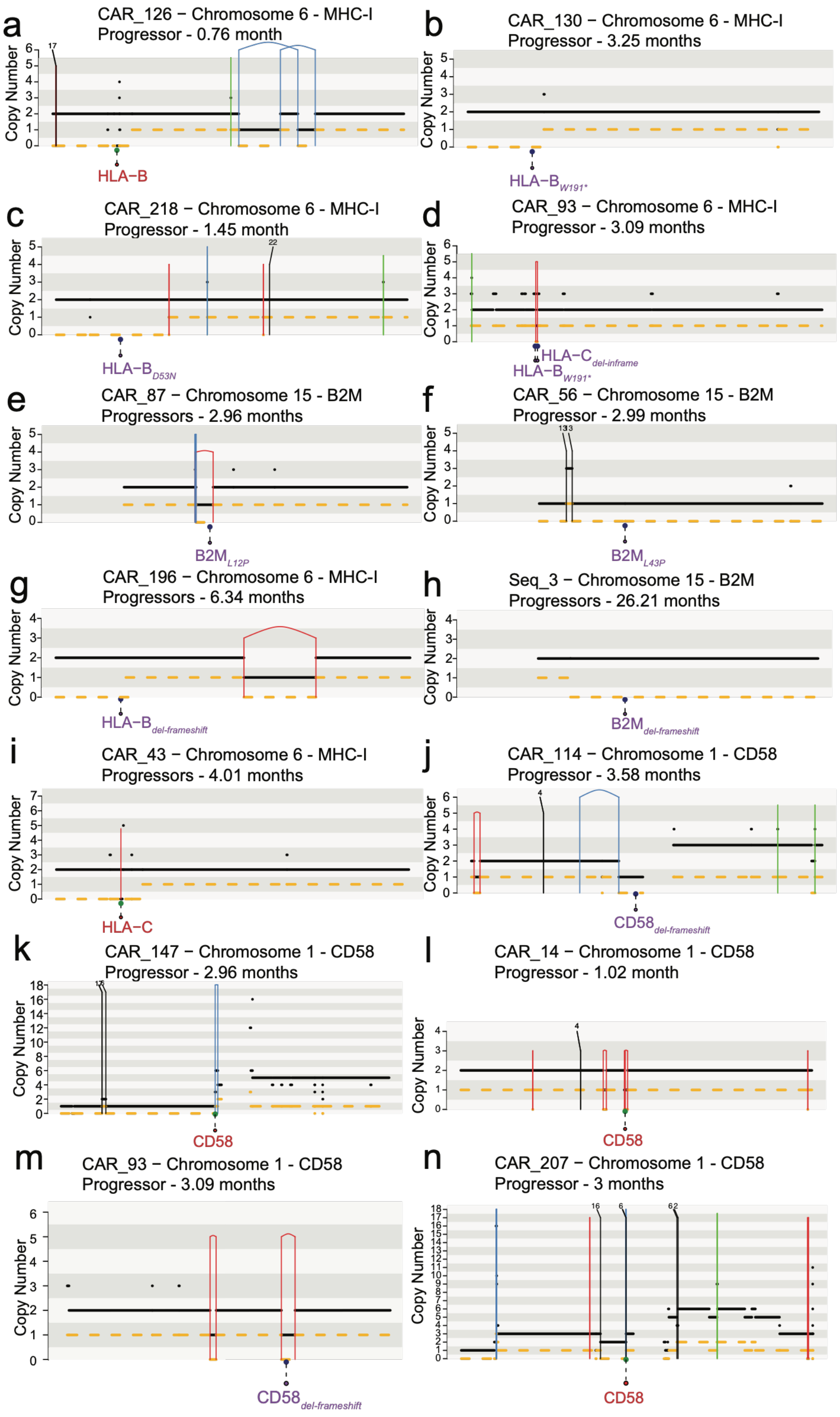

**Supplementary Fig. 8. Validation of B2m knockout and in vivo growth of A20-WT and A20-B2m-KO tumors. a.** Western blot confirming loss of B2m protein in A20-B2m-KO cells compared with A20-WT, with actin as loading control. **b.** Flow cytometry histograms showing surface expression of B2M, MHC-I, and CD19 in A20-WT versus A20-B2m-KO cells; unstained controls included. **c.** Bioluminescence imaging of tumor growth in mice bearing A20-WT or A20-B2m-KO tumors following injection of CAR-T cells. **d–e.** Tumor cell counts over time for A20-WT and A20-B2m-KO mice. **f.** Bar plot showing purified Balb/c NK cells were co-cultured with A20-WT (red) or A20-B2m-KO (blue) targets at the indicated effector:target (E:T) ratios; % lysis (mean ± s.e.m.) is shown. ns = not significant; *** = $p < 0.001$, **** = $p < 0.0001$ **g.** Representative flow cytometry plots of mCherry$^+$ A20 cells co-cultured with T cells.

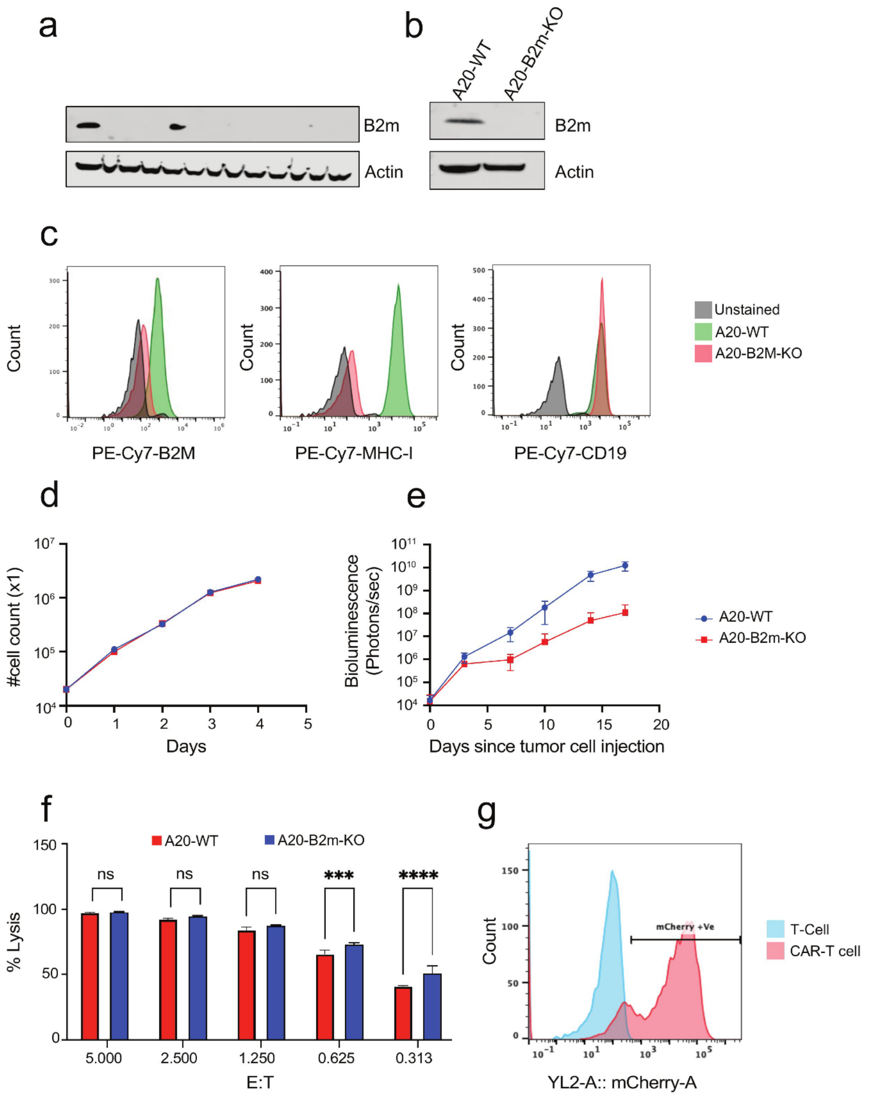

**Supplementary Fig. 9. Immune cell marker expression in the tumor microenvironment. a.** General UMAP plot of all samples, with the cell type information. **b.** Expression of T-cell markers (CD3E, CD4, CD8A). **c.** Proportion of cell type in WT and B2M-KO. **d.** Expression of the B-cell, monocyte and NK cell markers. **e.** UMAP plot of T cell subtypes. **f.** Bar plot comparing immune T cell subsets across conditions.

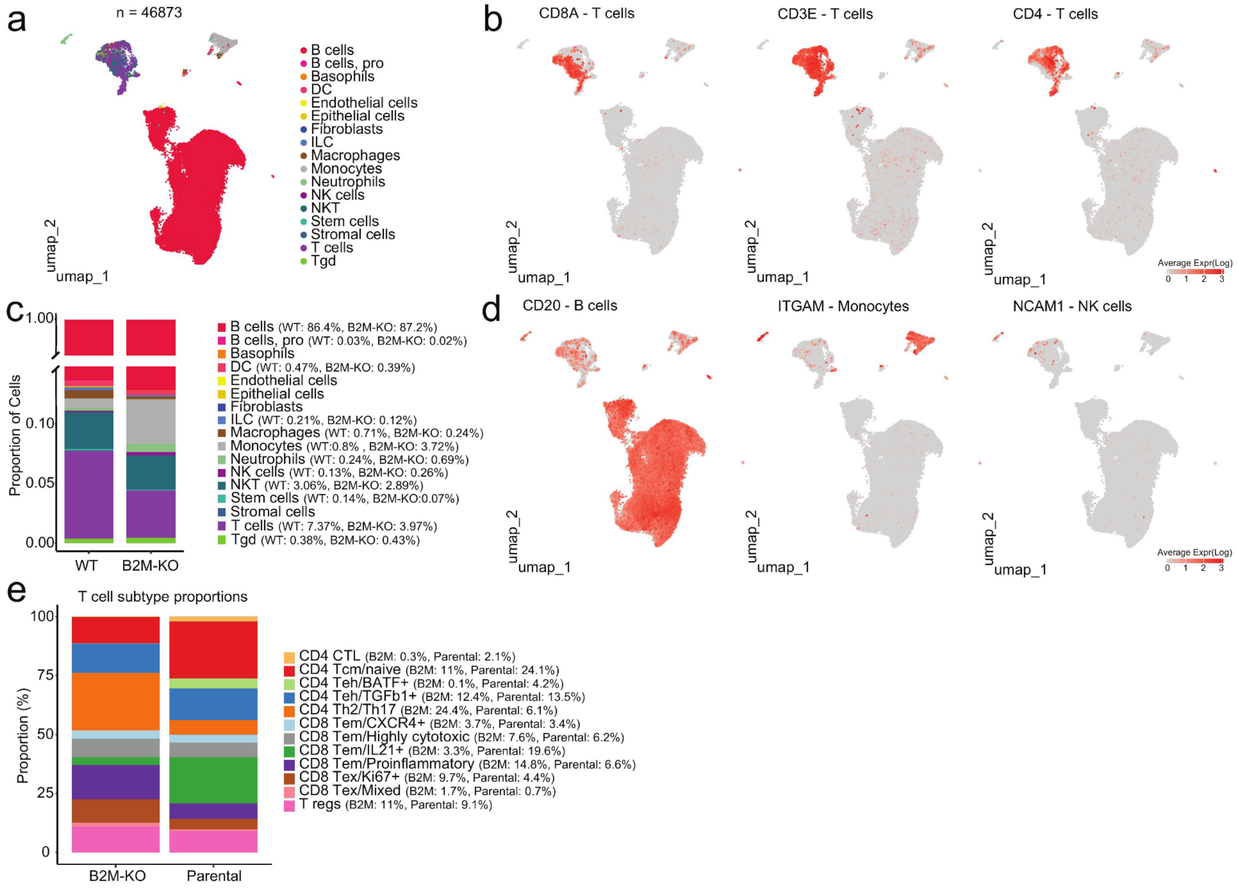

**Supplementary Fig. 10. Functional validation of B2m loss and CD8$^+$ T-cell dependency in CAR-T models. a.** Flow cytometry density plot confirming loss of B2M and MHC-I expression in BL3750 B2m-KO cells compared with controls. **b.** In vitro cytotoxicity assay showing percentage lysis of BL3750 target cells at different effector-to-target (E:T) ratios by CAR-T cells. **c.** Tumor growth curves over time (total flux, photons/sec) in WT versus CD8-KO hosts following CAR-T infusion. **d.** Flow cytometry plots showing CAR-T and T-cell populations in WT and CD8-KO hosts.

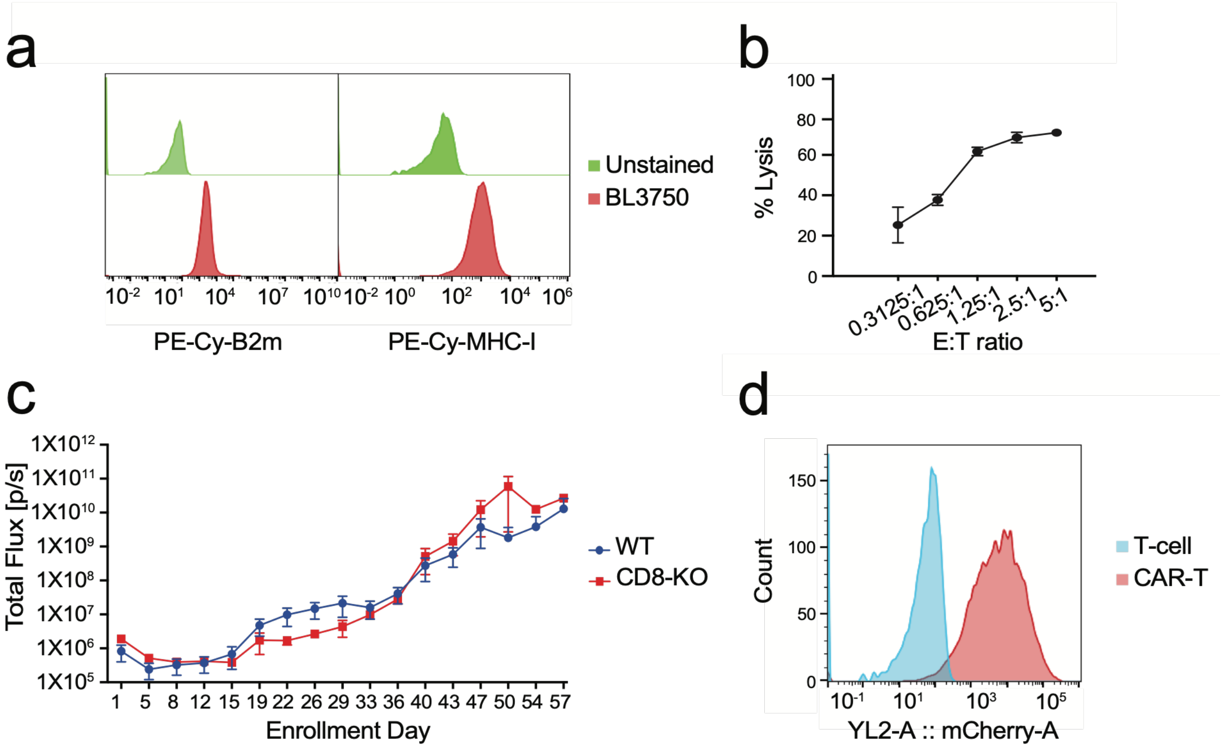

**Supplementary Fig. 11. Raw IVIS images.** Mice at D33, the last timepoint when all animals were still alive after engraftment and treatment. No tumors were found in CD8-WT hosts treated with CAR-T cells.

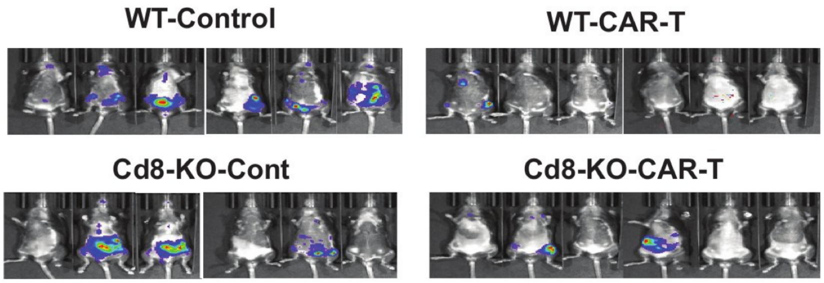

**Supplementary Fig.12. a.** Canonical marker expression used to annotate major cell populations. **b-c.** UMAPs showing cell distributions and annotated populations across experimental groups.

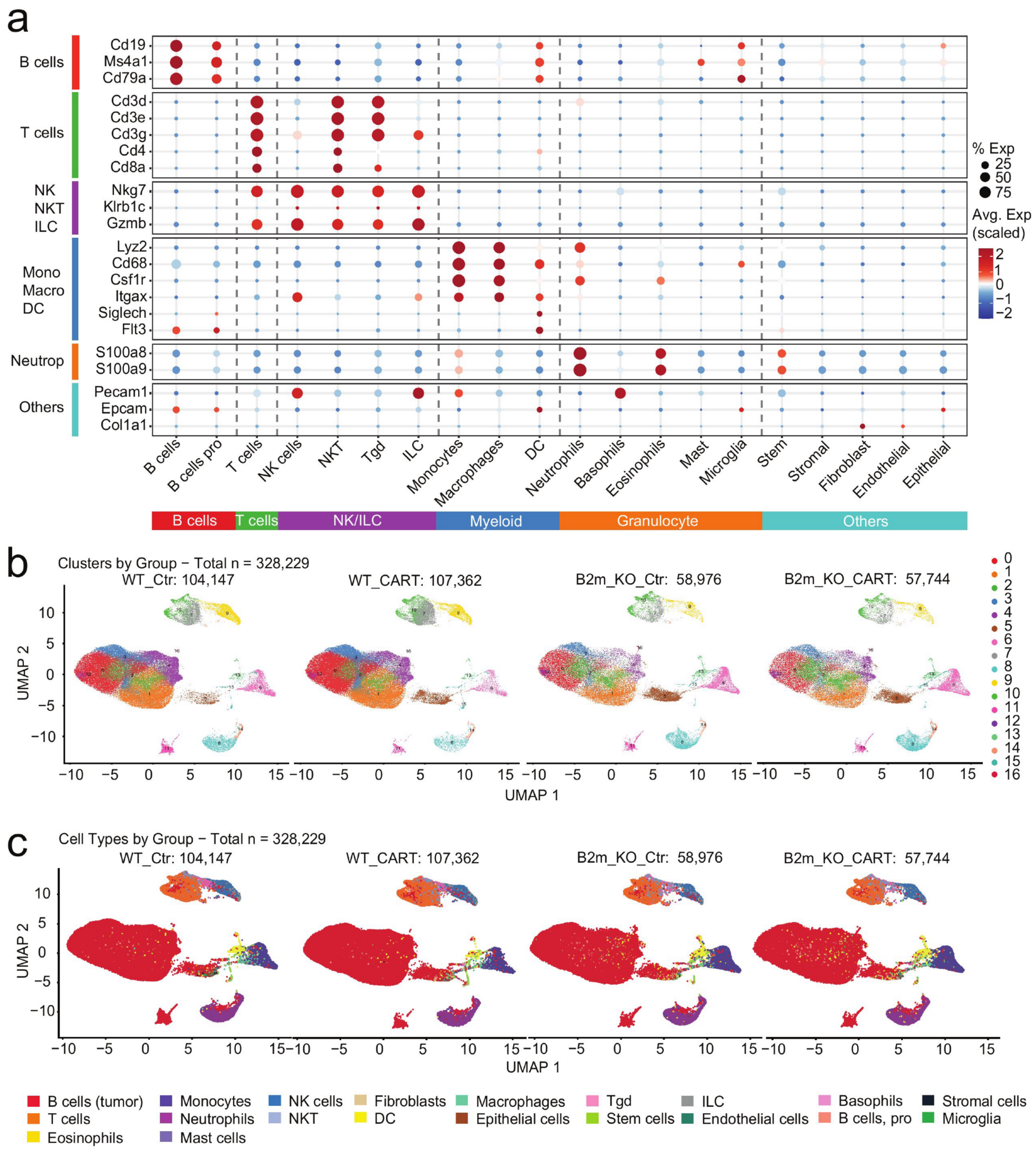

**Supplementary Fig.13.** Quantification of major tumor microenvironment populations using absolute cell counts; p-value computed using Wilcoxon test.

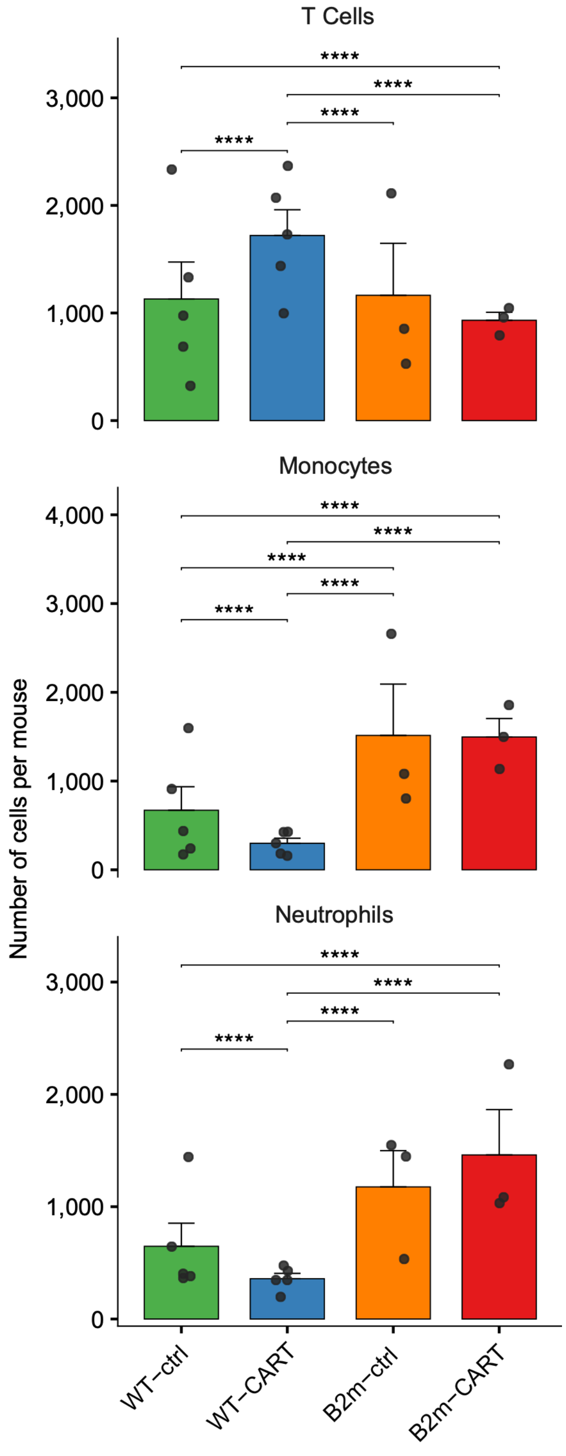

**Supplementary Fig.14. a.** Relative abundance of endogenous CD8 T-cell states across groups. **b.** Marker expression defining endogenous CD8. **c.** UMAP of re-clustered monocytes. **d.** Marker expression defining monocyte subpopulations. **e,f.** Hallmark pathway enrichment and differential gene expression analyses comparing monocytes from WTCAR-T and B2m-KO-CAR-T tumors.

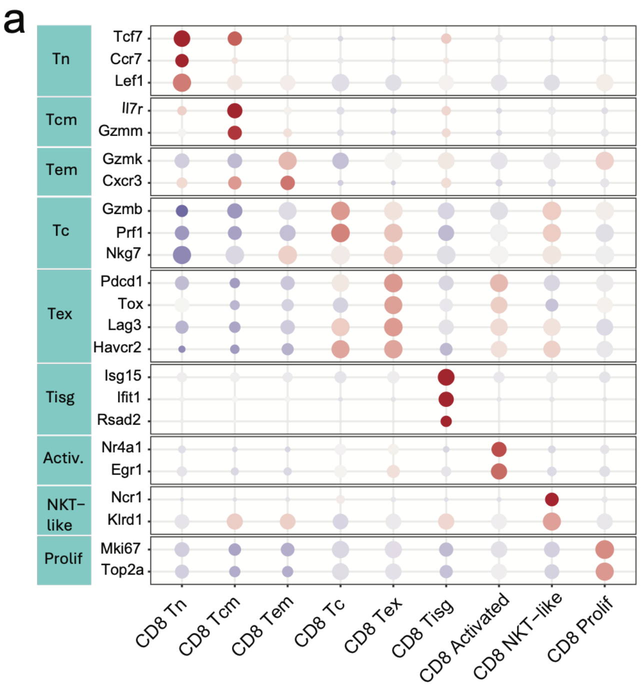


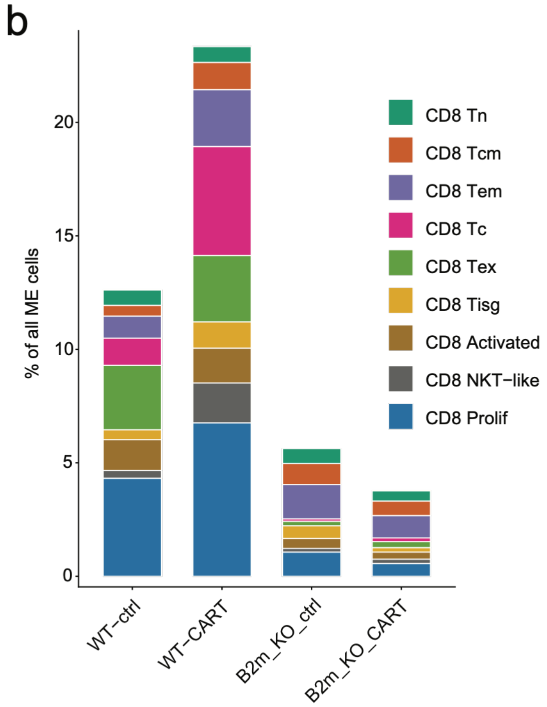

**Supplementary Fig.15. a.** Marker expression defining endogenous CD4. **b**. UMAP of endogenous CD4 T-cell subclusters. **c-d.** Relative abundance **(c)** and quantification **(d)** of endogenous CD4 T-cell states across groups.

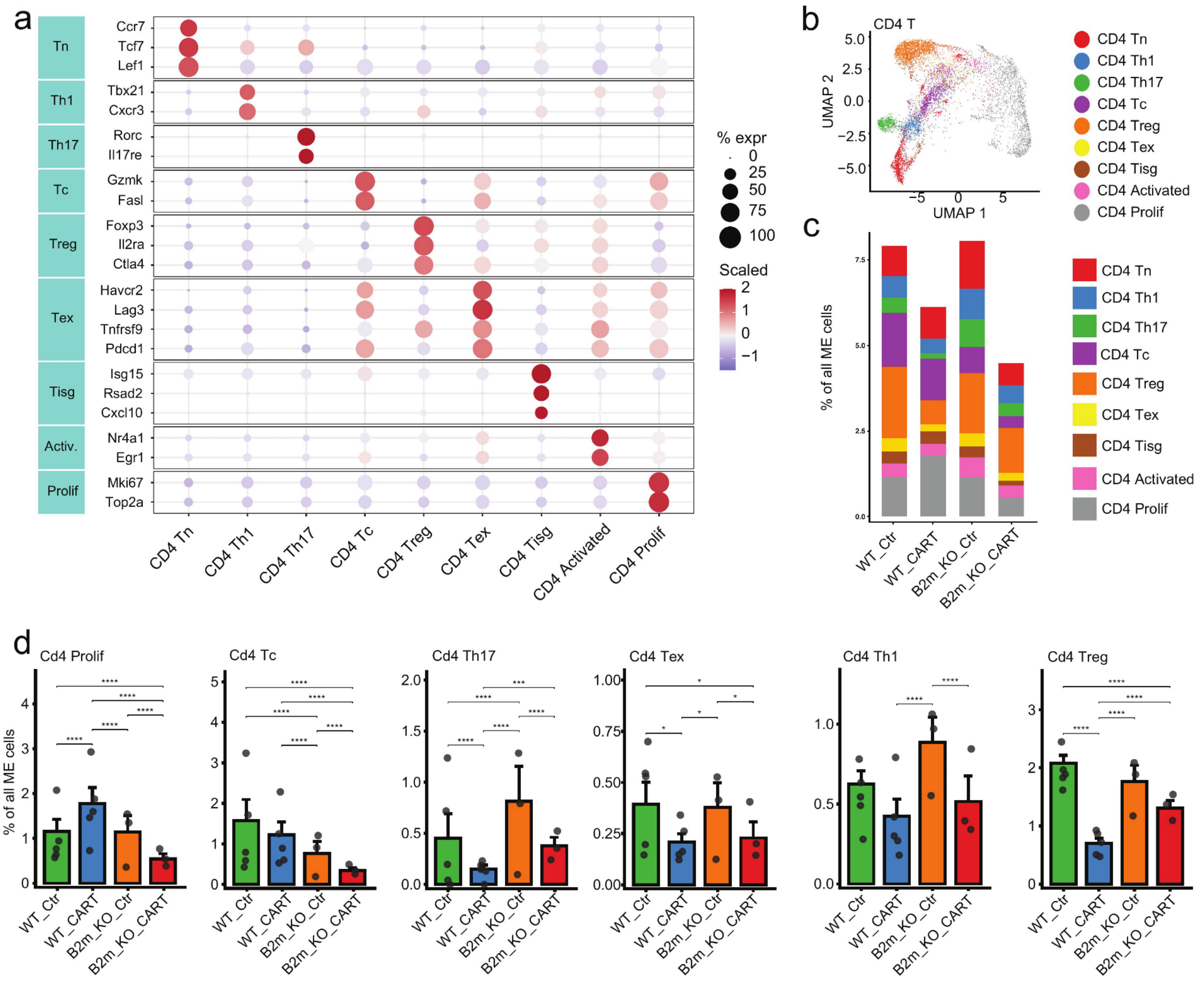

**Supplementary Fig.16. a.** UMAP of re-clustered monocytes. **b.** Marker expression defining monocyte subpopulations. **c,d.** Hallmark pathway enrichment and differential gene expression analyses comparing monocytes from WTCAR-T and B2m-KO-CAR-T tumors.

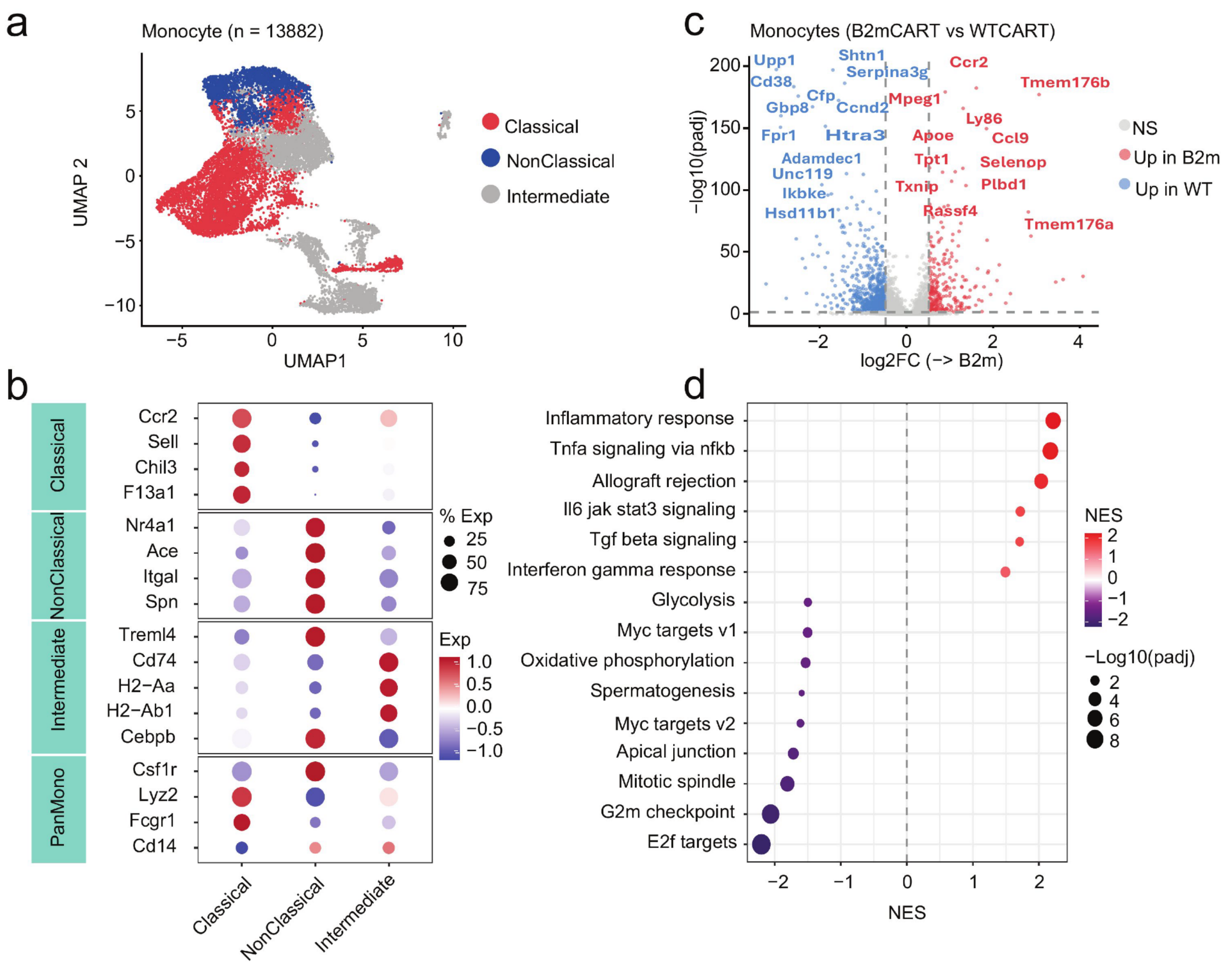

**Supplementary Fig. 17. a–b,** Circos plots depicting the global MultiNicheNet-predicted cell–cell communication landscape in CAR-T-treated WT **(a)** and *B2m*-KO **(b)** tumors. Ligand-expressing sender populations and receptor-expressing receiver populations are displayed around the circumference, with connecting ribbons representing prioritized ligand–receptor interactions. Ribbon color reflects the corresponding MultiNicheNet prioritization score. **c–e,** Prioritized ligand–receptor interactions for CAR-T cells→endogenous T cells **(c),** monocytes→endogenous T cells **(d),** and tumor B cells→endogenous T cells **(e)** in CAR-T-treated WT and *B2m*-KO tumors. Dot color indicates the MultiNicheNet prioritization score, and dot size indicates the fraction of cells expressing the corresponding ligand–receptor pair. Absence of a dot indicates that the interaction was not prioritized in that condition. **f–g,** Receiver-resolved MultiNicheNet analysis of communication from CD8$^{+}$ CAR-T cells→endogenous CD8$^{+}$ T cells **(f)** and CD4$^{+}$ CAR-T cells→endogenous CD8$^{+}$ T cells **(g)** in CAR-T-treated WT and *B2m*-KO tumors. Dot color indicates the MultiNicheNet prioritization score, and dot size indicates the fraction of cells expressing the corresponding ligand–receptor pair.

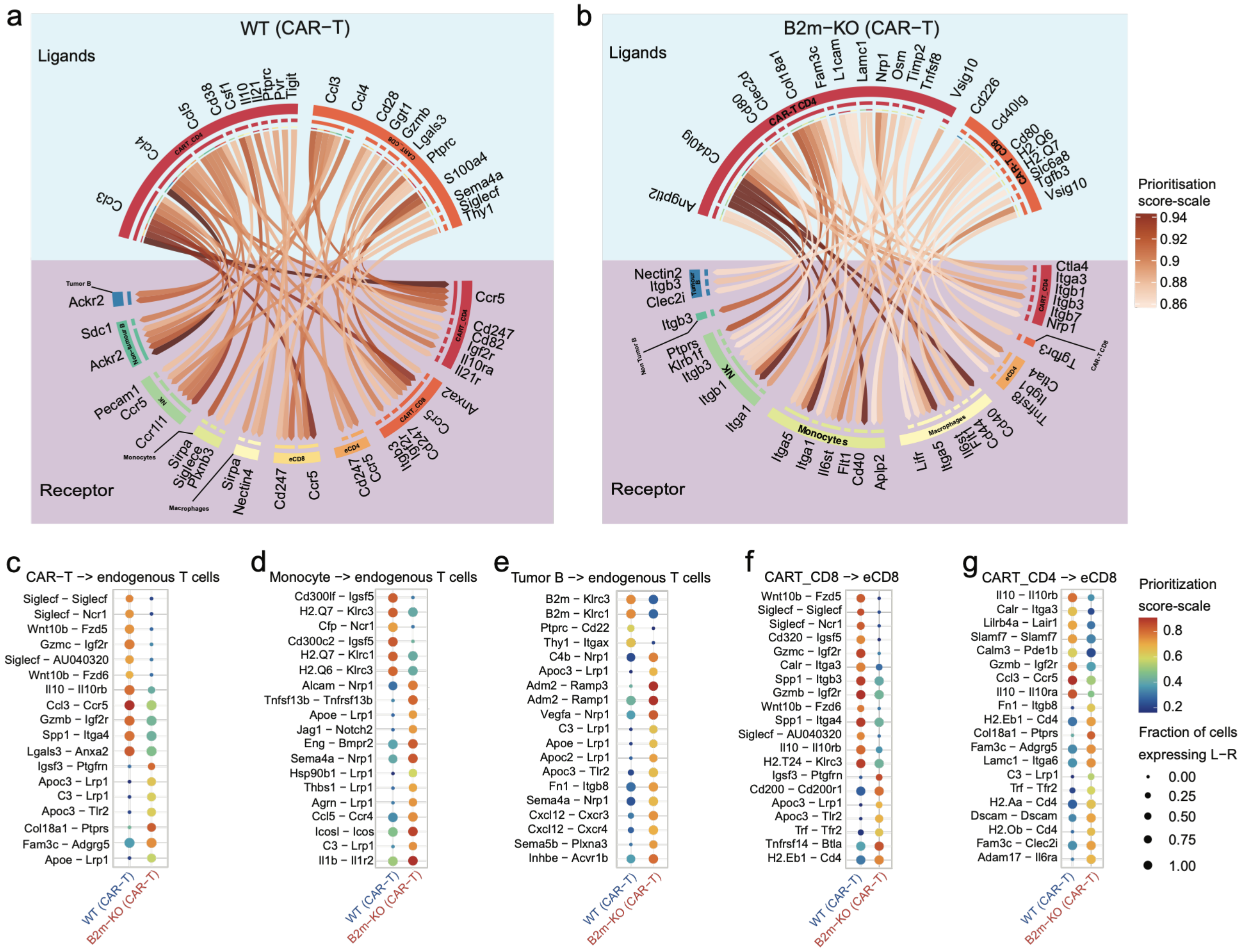

**Supplementary Fig. 18. IFNg ELISPOT from pre- or post-CAR-T infusion patient samples.** Representative images in triplicates of the IFNg ELISPOT from BL (left) or post-CAR-T infusion patient samples (right) cultured alone without stimulation (unstim), stimulated with PMA and ionomycin (PMA/Iono), or co-cultured with 3T3-null, 3T3-CD19 or autologous tumor cells (tumor). Tumor alone samples were included as controls.

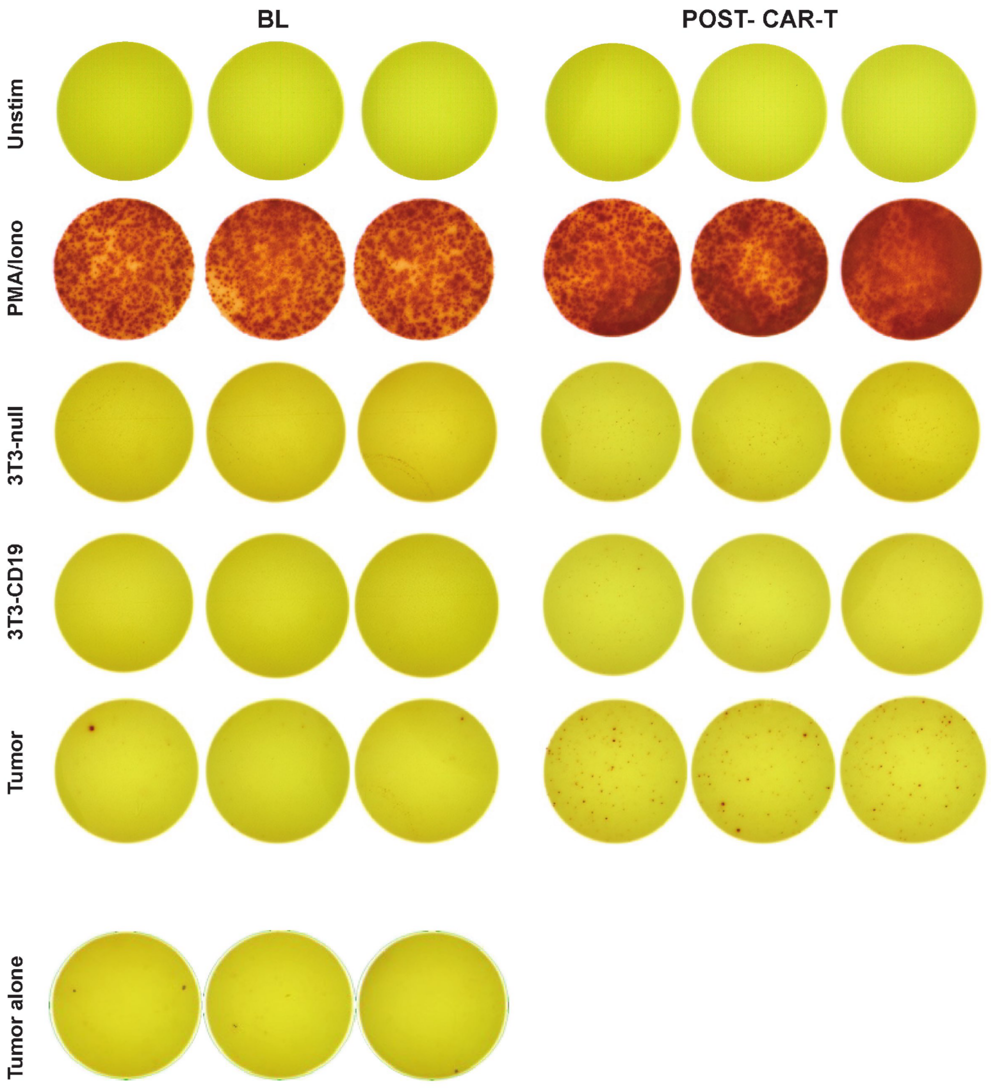

**Supplementary Fig. 19. IFNg ELISPOT from endogenous CD4 or CD8 samples. a.** Flow cytometry dot plots of representative mice from each experimental group demonstrating the enrichment of endogenous CD4 (eCD4, left) and CD8 (eCD8, right) T cells post FACS. **b.** Histograms from A20 WT, or A20 CD19KO and B2MKO target cells generated from parental A20 WT cells by CRISPR/Cas9, used as stimulation for ELISPOT demonstrating surface CD19 (left) or MHC-I (right) expression by flow cytometry. **c-d.** Representative images in duplicates or triplicates of the IFNg ELISPOT from endogenous CD4 (**c**) or CD8 (**d**) samples (right) from non-treated control mice (left) or CAR-T treated mice, cultured alone without stimulation (unstim), stimulated with PMA and ionomycin (PMA/Iono), or co-cultured with A20 WT, A20 CD19KO or A20 B2MKO (only for CD8 samples)

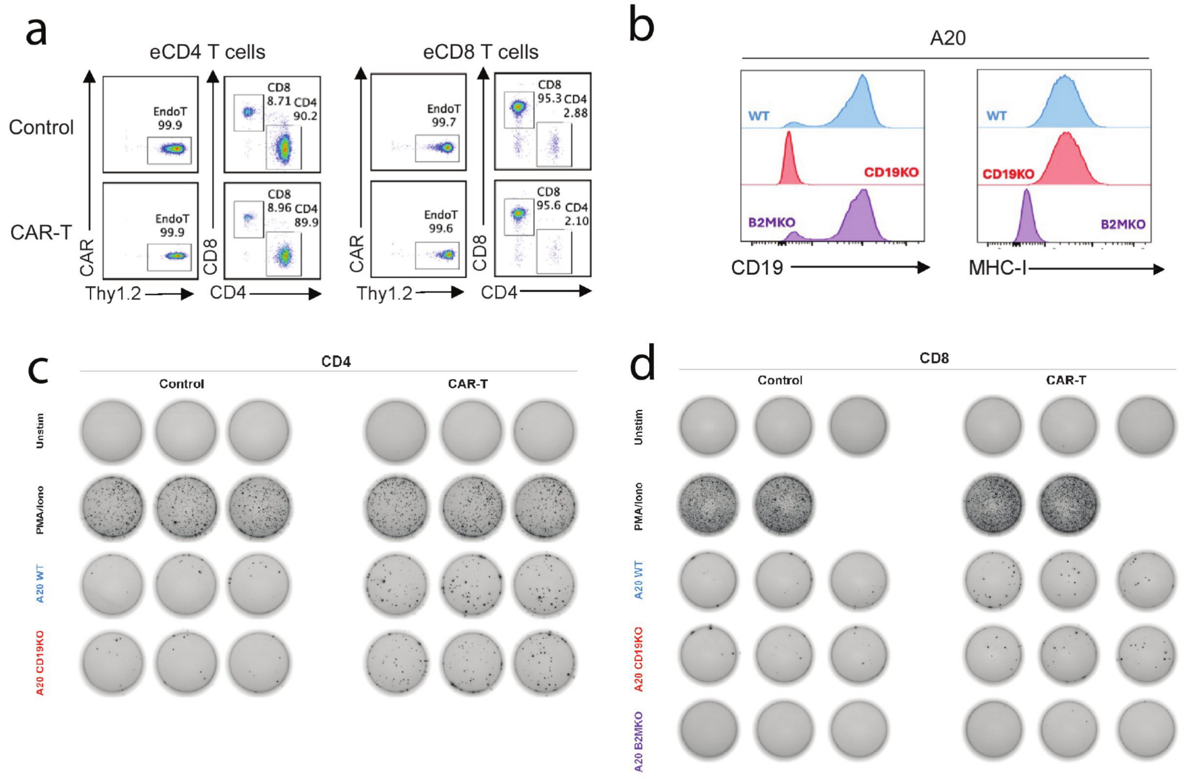
