## Supplementary Methods for "Lymphoma response to chimeric antigen receptor T cell therapy is driven by endogenous tumor-reactive CD8⁺ T cells"

### **Endogenous CD8⁺ T-cell are essential for anti-CD19 CART efficacy in large B-cell lymphoma**

**METHODS**

**Patient sample collection and preparation**

Patient samples included in this study were collected under an IRB-approved protocol, and all patients have provided written informed consent. LBCL patients received autologous CD19-targeted CAR-T cell therapy with axicabtagene ciloleucel. Surgical excisional tumor biopsies (tumor) were collected prior and after to CAR-T cell infusion, processed into viable cell suspensions and cryopreserved until use. Apheresis samples (Aph) were collected for CAR-T cell manufacture, and a small aliquot was cryopreserved for research purposes. Blood samples prior- (baseline, BL) or post-CAR-T cell infusion (post) were collected, and peripheral blood mononuclear cells (PBMCs) were enriched by Ficoll-Hypaque density gradient (StemCell, #18061) and cryopreserved until use.

**Analysis populations**

Sixty-one tumor biopsies with paired germline samples were obtained from 54 patients with r/r LBCL treated with CD19 CAR-T. All 54 patients contribute to the description of baseline characteristics (**Supplementary** **Table 2**). The population analyzed thereafter varies by analysis and is defined as follows; the flow of patients between analyses is summarized in **Supplementary Figure 1**.

Four patients (CAR_198, CAR_43, CAR_196 and Seq_3) achieved an initial complete remission followed by progression and were profiled only on a relapse biopsy. Because the progressing clone may be genomically distinct from the pretreatment tumor, these patients were excluded from all survival analyses, leaving 50 patients for overall survival (OS). One further patient (CAR_122) died within a week of infusion from CAR-19 toxicity with an unknown disease response; this patient was omitted from progression-free survival (PFS) analyses but retained in OS analyses, giving 49 patients for PFS. Three patients with primary refractoriness or limited response (CAR_26, CAR_218 and CAR_237) also had relapse-only biopsies, but their lack of response was attributable to pre-existing clonal alterations, and they were therefore retained.

Analyses relating PFS to pre-CART genomic features are based on 48 patients: CAR_16 was excluded because excessive copy-number fragmentation prevented assignment of the genomic-complexity signature. Multivariable Cox models additionally including baseline metabolic tumor volume and RNAseq samples are based on 41 patients, one having no baseline MTV measurement and 6 without expression data (and without evaluable neoantigen predictions; **Supplementary Table 3**).

| **Analysis** | **n** | **Relative to the preceding row** |
| --- | --- | --- |
| Baseline characteristics (Table 1) | **54** | All treated patients |
| Overall survival (OS) | **50** | − 4 transient CR with relapse-only biopsy |
| Progression-free survival (PFS) | **49** | − 1 early toxic death, response unknown (CAR_122) |
| PFS with pre-CART genomic features | **48** | − 1 without genomic-complexity call (CAR_16) |
| Multivariable Cox (incl. baseline MTV) | **47** | − 1 without baseline MTV |
| Neoantigen-burden analyses | **41** | − 6 without evaluable neoantigen predictions |

**Whole genome sequencing**

First, all DNA samples were evaluated for concentration by fluorometric Qubit assays (Thermo-Fisher) and for integrity by TapeStation (Agilent Technologies). Sequencing libraries were prepared using the TruSeq DNA PCR-free HT sample preparation kit from Illumina. Briefly, one ug of total genomic DNA was fragmented using the Covaris LE220 focus acoustic sonicator to a target size of 350bp. Blunt-end DNA fragments were generated and size selection performed with AMPure bead purification (Beckman Coulter). A-base tailing was performed on the 3’ blunt ends followed by adapter ligation and a bead-based clean-up of the libraries. Final library fragment size was evaluated on the TapeStation (Agilent Technologies) and final molarity quantification determined by qPCR with adapter specific primers (Kapa Biosystems) on a Roche Light Cycler. Libraries were normalized to 2.8nM and 24-samples pooled for sequencing on a S4-300 flow cell on the NovaSeq 6000. Paired-end 150bp reads were generated to yield an average depth of 30x per sample. FASTQ files were generated using the Illumina BCL2FASTQ algorithm and used for downstream processing.

**Whole genome sequencing analysis**

The MGP1000 pipeline (<https://github.com/pblaney/mgp1000>) was used to align tumor and normal paired samples against the GRCh38 reference genome using the Burrows–Wheeler Aligner. All samples were uniformly analyzed by the following bioinformatic tools: single nucleotide variants (SNVs) and small insertion-deletions (INDELs) were identified by integrating MuTect2, Strelka2 and VarScan, selecting the variants called by at least 3 callers and applying additional filters to remove artifactual variants; copy number analysis (CNVs) and tumor purity (i.e., cancer cell fraction) were evaluated using ASCAT (<https://github.com/VanLoo-lab/ascat>); structural variants (SVs) were defined by Manta (<https://github.com/Illumina/manta>), SvABA (<https://github.com/walaj/svaba>), DELLY2 (<https://github.com/dellytools/delly>) and IgCaller algorithms^1^. Complex events (i.e., templated insertions, chromothripsis, and chromoplexy) were defined after manual inspection, as described previously^2^.

**Mutational Signatures**

Mutational signatures were analyzed across all whole genomes. To estimate the activity of mutational signatures, we first used a three-step approach of de novo extraction, assignment, and fitting^3,4^. For the first step, we ran SigProfiler for SBS signatures. All extracted signatures were then compared with the latest Catalogue of Somatic Mutations in Cancer (<https://cancer.sanger.ac.uk/cosmic/signatures/SBS>; COSMIC) to identify the known mutational processes active in the cohort. Finally, we applied mmsig (<https://github.com/UM-Myeloma-Genomics/mmsig>)^4^, a fitting algorithm, to confirm the presence and estimate the contribution of each mutational signature in each sample guided by the catalog of signatures extracted for each individual sample adding the chemotherapy SBS-MM1 (SBS99) and platinum signatures for patients who received melphalan^5^.

**RNA sequencing analysis**

The RNA sequencing (RNA-seq) libraries were prepared with the NuGen RNA-Seq Multiplex System (Tecan US) as previously described^6^. The libraries were sequenced on the Illumina NextSeq 500 system with a 75 base paired end run at 80 to 100 million read pairs per sample. RNA-seq reads were mapped to the reference human genome (GRCh38) using the STAR algorithm ([https://github.com/alexdobin/STAR](https://nam10.safelinks.protection.outlook.com/?url=https%3A%2F%2Fgithub.com%2Falexdobin%2FSTAR&data=04%7C01%7Ccac451%40med.miami.edu%7Cc48117faff1a4efda59f08d9699e7bae%7C2a144b72f23942d48c0e6f0f17c48e33%7C0%7C0%7C637656951661140917%7CUnknown%7CTWFpbGZsb3d8eyJWIjoiMC4wLjAwMDAiLCJQIjoiV2luMzIiLCJBTiI6Ik1haWwiLCJXVCI6Mn0%3D%7C1000&sdata=Ml4%2F%2BNgNByzFulivz7iAQmqEtXM8%2Fjsy1xZ1gauHV8E%3D&reserved=0)), and the parameters were set to count read numbers per gene while mapping.

**Genomic complexity signature extraction and assignment**

To provide an estimation of the level of genomic complexity in each sample, we extracted de novo copy number variation (CNV) and structural variant (SV) signatures using the Hierarchical Dirichlet Process (HDP)^7^, integrating CNV and SV profiles from whole-genome sequencing data. To improve resolution and statistical power, we combined our r/r LBCL cohort, with 51 additional newly diagnosed LBCL cases from the PCAWG dataset^8^. After excluding samples with excessive copy-number fragmentation (CAR_16) or high BRCA-SV burden (DO52686), HDP identified 7 CNV signatures and 4 SV signatures Signatures were evaluated, using the PCAWG dataset, for association with known complex genomic events, including chromothripsis, pyrgo, rigma, breakage-fusion-bridge (BFB), and double minutes. Next, we selected the signatures which were enriched in patients with these alterations. Of these, CNV.Sig1, CNV.Sig5, CNV.Sig6, SV.Sig2, and SV.Sig4 were significantly enriched (p < 0.05, Wilcoxon test; **Supplementary Table 10**). These were defined as the complex CNV-Sig and the complex SV-Sig, both characterized by complex localized and clustered rearrangements and multiple large and focal aneuploidies. We then applied these complexity-associated signatures to our CD19 CAR-T cohort and observed consistent associations, supporting the robustness and cross-cohort reproducibility of the signatures. To delineate the overall genomic complexity signature, we identified the smallest cutoff for each complex signature (CNV and SV) using a log-rank test and assigned “complex genome” status when at least one of these CNV or SV signatures was present. This approach yielded a composite genomic complexity signature, which was subsequently applied to the CD19 CAR-T cohort for downstream analyses.

**Clonal architecture inference**

To reconstruct tumor phylogenies and infer clonal evolution, we used the MGP1000 pipeline to generate somatic variant calls from whole-genome sequencing (BAM) files. Somatic single-nucleotide variants (SNVs) and copy-number variants (CNVs) were processed with DpClust (<https://github.com/Wedge-lab/dpclust>), which integrates purity estimates to derive cancer cell fractions (CCFs) for each mutation. Mutations were clustered into subclonal populations based on their CCFs, and clusters were used to build patient-specific phylogenetic trees. These trees allowed annotation of clonal versus subclonal events and provided a framework to assess evolutionary trajectories and branching patterns within each tumor.

**Neoantigen prediction and immune-evasion analysis**

Somatic variants derived from the MGP1000 pipeline were analyzed to assess MHC-I status and predict neoantigen presentation. MHC class I alleles were inferred using Polysolver^9^, and allele-specific copy number loss was evaluated with LOHHLA (<https://github.com/mskcc/lohhla>). To generate a confident list of MHC-I biallelic loss events, we combined the LOHHLA output with somatic variants (CNVs, SVs and SNVs/INDELs). Biallelic loss were identify with homozygous deletion or deletion plus non-synonymous mutation with clonal impact on protein function. Predicted mutations were required to have CCF ≥ 0.9 and a functional effect (PolyPhen-2, SIFT, CADD, or PhyloP).

A functional effect is defined when:

| **Algorithm** | **Condition** |
| --- | --- |
| PolyPhen-2 | ≥ PROBABLY damaging |
| SIFT | = DELETERIUS |
| CADD | = HIGH LIKELY deleterius |
| PhyloP | = HIGHLY CONSERVED |

A variant is reported with functional impact when:

| **Type of mutation** | **Missense** |
| --- | --- |
| Missense | Clonal + at least 2 algorithms |
| Nonsense | Clonal + at least 1 algorithm |
| Frameshift | Clonal |

The same approach was applied to assess CD58 biallelic inactivation. To create a neoantigen confident list, SNSv/INDELs were analyzed using pVACseq to predict candidate neoantigens, corrected for the MHC-I allele loss (neoantigens presented by the MHC-I loss allele were removed from the analysis), and expression was confirmed by RNA-seq to exclude non-expressed peptides.

**Computational model**

We developed a multi-predator (endogenous T and CAR) multi-prey (tumor clones) consumer-resource model^10^, based on our previous work^11,12^. Where the interaction between CAR-T and intra-tumor endogenous CD8 T-cells (eCD8) comes in the form of cooperative hunting^13^, wherein higher abundances of predators increase the detection and killing rates of the tumor cell population. We run several simulations, where the parameters were found by fitting the model to clinical data provided in. Each simulation begins with a small, homogenous tumor that has an initial immunogenicity, CD19 and MHC-I expression levels, all of which range between 0 and 1. The tumor generates mutants as the cells within it divide, and each new mutant acquires a novel neoantigen with a randomly assigned immunogenicity. Parental neoantigens are inherited, meaning that neoantigens accumulate over time, and can be measured at the time of CAR-T infusion. The pre-treatment neoantigen burden of each simulated patient is thus shaped by selection against immunogenicity, which can be mitigated via lower MHC-I expression levels. Following 90 simulated days of tumor growth, treatment is administered. Lymphodepletion reduces the number of eCD8, and CAR-T are then added to the simulation. There is an initial proliferating subpopulation of CAR T cells that expand following infusion, but the size of this population decreases over time, and eventually CAR-T cease to expand. Throughout treatment, all CAR T are capable of attacking tumor cells at a rate proportional to the tumor’s CD19 expression levels. While CAR T killing is MHC-I independent, and eCD8 killing is CD19 independent, cooperative hunting means that the presence of each T-cell population enhances overall tumor kill from both T-cell populations. Following infusion, the simulation is run for an additional simulated 190 days, or until the tumor is eliminated, whichever comes first. To account for patient variability, we perform a parameter sweep over different combinations of initial immunogenicity, CD19 expression, and HLA expression. While most patients’ HLA levels are likely closer to the extremes, e.g. HLA- or HLA+, this sweep makes it possible to fully explore the impact of HLA, even in the theoretical/rare range of moderate HLA expression. The Latin Hypercube sampling strategy^14^, was used to select 500 combinations that sufficiently cover this 3D parameter space. Pre-treatment neoantigen burden was determined by counting the number of cells that carry each neoantigen, which requires traversing through each clone’s descendants to count all cells that inherited the neoantigen. To account for limitations in sequencing depth, we select a mutation detection threshold that yields neoantigen counts similar to those in the observed data. It is worth noting that this neoantigen burden data comes from a different cohort than the one used to parameterize the model. The distribution of pre-treatment neoantigen burden can then be compared between the simulated tumors that progressed and those that did not.

**Cell lines**

3T3 and A20 cells were obtained from American Type Culture Collection (ATCC) and cultured as recommended, in complete RPMI: RPMI 1640 (Gibco) supplemented with 10% fetal bovine serum (FBS) and MycoZap-Plus-CL (Lonza, VZA-2012). For A20 cells, High glucose ATCC modificated-RPMI was used and supplemented as described above, plus 2-mercaptoethanol 0.5mM (ThermoFisher, #31350010). To generate 3T3 cells overexpressing human CD19 (3T3-CD19) 3T3 cells were transduced in a retronectin-coated plate with a gammaretrovirus containing a DNA plasmid encoding the human CD19 protein, spun for 1h at 2000g 32C and incubated at 37C 5%CO_2_. A week after, CD19+ cells were enriched by FACS sort. Green fluorescent protein (GFP) and firefly luciferase (FFL) were incorporated into A20 by transduction with a gammaretroviral vector encoding both GFP and FFL, following the steps mentioned above. Cells were FACS sorted based on GFP+ expression. CD19 knock out (CD19KO) and beta-2microglobulin knock out (B2MKO) A20 cells were generated from parental A20 wild type (WT) GFP FFL cells by CRISPR/Cas9. Respective guides and reagents were pre-designed and obtained from IDT. KO cells were purified by FACS sort. **CRISPR/Cas9-mediated knockout of B2*m* using parental A20-WT cells** was performed via transient co-nucleofection (Lonza Nucleofector Kit V, VCA-1003) using eSpCas9(1.1)-T2A-GFP-GlntRNA and eSpCas9(1.1)-T2A-mCherry-GlntRNA cloned with sgRNAs targeting exon 2 of B2*m* (guide 1: CACGCCACCCACCGGAGAAT; guide 2: TCACGCCACCCACCGGAGAA) for 24 h, followed by FACS sorting of double-positive (GFP/mCherry) single cells in 96-well plates. Single clones were expanded, and **B2*m*-KO** was assessed by Western blot (WB) and confirmed by loss of surface β2-microglobulin expression via flow cytometry along with MHC-I and *CD19* expression levels compared to A20-WT. The proliferation differences between WT and B2m-KO cells were determined for 72 hr. For in-vivo, both A20-WT and A20-B2*m*-KO cells were transduced with the lentiviral construct pCDH-MSCV-Luciferase-EF1-GFP at an MOI of 4. Three days after transduction, GFP-positive cells were isolated by cell sorting. All cells passed authentication and tested negative for mycoplasma (ATCC, #301012K) prior to starting the experiments.

**Mice**

Balb/cJ wild type (WT) and CBy.PL(B6)-Thy1a/ScrJ Balb/c (Thy1.1+) female mice were purchased from Jackson Laboratories (JAX #651 and #5443, respectively). Balb/c-WT for A20-WT/ B2*m*-KO experiment, C57BL/6J (WT) and B6.129S2-Cd8atm1Mak/J (CD8-KO) mice were purchased from Jackson Laboratories (JAX # 000651, JAX # 000664 and JAX # 002665 respectively). All experimental protocols were approved by the Institutional Animal Care and Use Committee (IACUC).

**Gamma-retroviral plasmid editing and virus production**

The SFG retroviral backbone was modified to include (1) human CD19 protein, (2) GFP FFL or (3) Ametrine-P2A-mCD19CAR (fluorescent protein ametrine, followed by a P2A protein and an anti-mouse CD19-targeted single-chain variable fragment (ScFv) with a CD8α transmembrane and hinge domain, CD28 costimulatory domain and CD3ζ. All plasmids were synthesized externally (Genewiz) and verified by restriction enzyme digest. Construct were calcium phosphate transfected into H29 packaging cells. Retroviral supernatants of transfected H29 were harvested and used to transduce stable producers PhoenixE cells. Retroviral supernatant of producer cells was harvested, 0.45 µm filtered, and used to transduce target cells (1 and 2) or T cells (3). Mouse SFG-m1928z-mCherry was gifted by Dr. Marco Davila. Retroviral particles carrying the SFG-m1928z-mCherry construct was produced by co-transfecting 293T cells (ATCC CRL-3216) with the transfer plasmid and the pCL-eco packaging plasmid at a 3:1 ratio. Approximately 6 × 10⁶ 293T cells were plated in 150-mm culture dishes one day before transfection. Construct were transfected using Lipofectamine 3000 (Thermo Fisher, L3000001) according to the manufacturer’s guidelines. After six hours, the transfection medium was gently replaced with fresh DMEM supplemented with 10% FBS and 100 IU/mL penicillin/streptomycin. Viral supernatants were collected 48 hours later, filtered and concentrated overnight with the Retro-X Concentrator (Takara, 631456) to a final volume of 100 µL per aliquot, and stored at −80 °C until needed.

**Murine CAR-T Cell Production via Retroviral Transduction**

SFG-m1928z-mCherry CAR-T; on day-0, 8–12-week-old either C57BL/6 or Balb/c Thy1.1⁺ mice were humanely euthanized, and spleens were harvested into cold 2% FBS in PBS. Spleens were gently dissociated through a 70 µm cell strainer to obtain single-cell suspensions, which were centrifuged at 400 X g for 10 min at 4^0^ C and washed twice with 2%FBS in PBS. CD3⁺ T cells were isolated using the EasySep™ Mouse T Cell Isolation Kit (StemCell # 19851) and resuspended at 1 × 10⁶ cells/mL in complete RPMI 1640 medium supplemented with 10% fetal bovine serum (FBS), 1 mM sodium pyruvate, 0.1 mM non-essential amino acids, 10 mM HEPES, 50 µM β-mercaptoethanol, and 100 U/mL penicillin/streptomycin. Cells were activated overnight with Dynabeads™ Mouse T-Activator CD3/CD28 (Invitrogen #11452D) at a 1:1 bead-to-cell ratio in the presence of recombinant mouse 50 IU/mL of IL-2 (R&D systems # 202-IL-050/CF) to promote proliferation. On day 1, non-treated 6-well plates were coated with 20 µg/mL Retronectin (Takara # T202) overnight at 4 °C, blocked with 2% bovine serum albumin (BSA) for 30 min, washed with PBS, and loaded with concentrated retroviral supernatant (MOI of 4) in 1.4 ml of RPMI media. Plates were centrifuged at 2,000 × g for 2 h at 32 °C to pre-bind virus. After pre-binding removed the viral supernatant and overnight activated mouse T cells were collected by centrifuging at 400 X g for 5 min and plated onto pre-bind viral wells at 2-3 × 10⁶ cells/well, followed by centrifugation at 800 × g for 30 min at 32 °C (spinoculation). Cells were incubated overnight at 37 °C with 5% CO₂. Beginning on day 2, cultures were expanded in complete RPMI supplemented with 50 IU/mL IL-2, refreshing medium to support expansion and viability. Transduced CD3⁺ T cells were monitored for gene expression by flow cytometry (mCherry^+^) and mCherry^+^ cells were enriched by FACS sort on day 4. Flow sorted CAR-T cells were incubated overnight at 37 °C with 5% CO₂ complete RPMI supplemented with 50 IU/mL IL-2. Overnight (Day-5) expanded pure CAR-T were used for downstream experiments and adoptive transfer.

**B cell lymphoma mouse model and CAR-T cell manufacture**

Thy1.1+ CAR-T cell manufacture

On day 1, 6-10-week-old Thy1.1+ mice were humanly euthanized, and spleens were collected and processed. Red blood cells were lysed using ammonium-chloride-potassium buffer (ThermoFisher) for 10min at room temperature (RT) and washed with phosphate buffer saline (PBS, Gibco). T cells were isolated by negative selection using a magnetic mouse T cell isolation kit (StemCell, #19851), activated with mouse T cell dynabeads at 1:1 ratio (Invitrogen, #11452D) and cultured overnight in complete mouse RPMI (RPM 1640 supplemented with 10% FBS, MycoZap-plus-PR (Lonza, #VZA-2022), minimum essential medium with non-essential amino acids (MEM NEAA, Gibco, #11140-050), 1mM Sodium Pyruvate (Gibco, #11360-070), 55μM β-Mercaptoethanol (#21985-023), 10mM HEPES (Gibco, #15630-080) in the presence of 100IU/mL of IL-2 (R&D Systems, #202-IL-050). On days 2 and 3, T cells were transduced in retronectin-coated 6-well plates with Ametrine+anti-mouse CD19 CAR-encoding gammaretroviral vectors and spun 1h at 2000g 37C and incubated overnight at 37C 5%CO_2_. On day 4, fresh complete mouse RPMI supplemented with IL-2 (100IU/mL) is added to the T cells, and an aliquot of T cells are stained for flow cytometry with an anti-G4S PE antibody (Cell Signalling Technology, #38907L) and live/dead staining for CAR transduction efficiency evaluation. On day 5, dynabeads are magnetically removed from the cells, T cells washed twice with PBS and resuspended at 3x10^6^ cells/200uL sterile PBS for adoptive transfer.

Mouse B cell lymphoma model

A20-WT/ B2*m*-KO lymphoma model: 8 to 12-weeks old Balb/c-WT mice were used for all experiments. On the morning of tumor cell injection mice received a single 3.5 *Gy* total-body irradiation. After irradiation (defined as Day 0) mice were injected via tail vein with 2.5 × 10^5^ tumor cells per mouse of either A20-WT-Luc or A20-B2*m*-KO-Luc cells. Mice were divided into four groups (n = 5 per group, total N = 20): A20-WT control, A20-WT-CAR-T, A20-B2*m*-KO-control, and A20-B2*m*-KO-CAR-T. For treatment groups, 1X10^6^ cells/mice of CAR-T cells (5 days post-activation/culture) were delivered intravenously via tail vein on day-5 after tumor cell injection. Tumor burden was assessed twice weekly using IVIS imaging for luciferase-expressing tumors and by clinical observation, while overall survival was monitored, with mice euthanized upon reaching humane endpoints. For single-cell RNA-seq analysis, irradiated Balb/c-WT mice were injected via tail vein with 1 × 10^6^ tumor cells per mouse of either A20-WT-Luc or A20-B2*m*-KO-Luc cells (n=5). CD-KO model: C57BL/6J (WT) and B6.129S2-Cd8atm1Mak/J (CD8-KO) mice (8–12 weeks old) were used. On the day of tumor challenge, mice received 3.5 Gy total-body irradiation, followed by intravenous injection of 5 × 10⁵ BL3750-Luc cells (a CD19⁺ B-cell lymphoma derived from *c-Myc* transgenic C57BL/6). Mice were randomized to untreated or CAR-T–treated cohorts, with treatment groups receiving 5-day–activated and cultured pure-CAR-T cells (2 × 10⁶ cells/ mice). Tumor burden was assessed twice weekly using an IVIS Spectrum (Revvity) and by clinical observation, while overall survival was monitored, with mice euthanized upon reaching humane endpoints.

Six to ten-week-old WT Balb/c mice were intravenously (iv) injected with 1×10^6^ A20 WT-GFP/FFluc at day 0, and 3×10^6^ CD19-targeted Thy1.1+ CAR T cells were intravenously injected at day 7. A group of non-treated mice was kept as controls. Tumors were measured using an IVIS 200 In Vivo Imaging System (PerkinElmer) for bioluminescence imaging (BLI) weekly. Mice were monitored for illness daily and sacrificed when there was evidence of leukemia progression, such as decreased activity, hunched posture, or ruffled coat. Two weeks after tumor challenge mouse were humanely euthanized, and spleens were collected and processed. Red blood cells were lysed and T cells isolated as described above. Samples were incubated with FcR block antibody (BD, #553141) and stained with anti-Thy1.1 PE (Biolegend, #202524), anti-G4S PE, anti-Thy1.2 APC (Biolegend, #105312) and anti-CD4 PE-Cy7 (Biolegend, #116016) antibodies for 30min at 4C. Samples were washed and resuspended in MACS Quant Tyto buffer (Miltenyi, #130-107-206) with Sytox Green (ThermoFisher, #S7020) for live/dead discrimination. Samples were filtered and sorted in high speed Tyto cartridge (Miltenyi, #130-121-549) using a MACS Tyto cell sorter (Miltenyi). Endogenous CD8 T cells (eCD8) were sorted as SytoxGreen- Thy1.1/G4S- Ametrine- Thy1.2+ CD4-. Negative fractions were collected, filtered and re-sorted on a new cartridge to isolate endogenous CD4 T cells (eCD4) as: SytoxGreen- Thy1.1/G4S- Ametrine- Thy1.2+ CD4+. Both positive fractions (eCD8 and eCD4) were collected and incubated overnight at 37C 5%CO_2_ in a humidified incubator. Pre and post sort aliquots were taken and stained for 30min with anti-CD8 BV605 antibody (Biolegend, #100744), washed and acquired in a BD LSR II flow cytometer. Data was analyzed using FlowJo VX (Tree Star).

Tumor rechallenge studies

BALB/c mice were inoculated with 5×10^5^ A20 WT (CD19^+^) GFP/FFL lymphoma cells and, following confirmation of tumor engraftment and normalization of tumor burden, were either treated with 3×10^6^ CD19-directed CAR-T cells or left untreated as controls. Mice achieving durable complete responses after CAR-T cell therapy were rechallenged 100 days after initial tumor inoculation with GFP/FFL CD19KO A20 lymphoma cells, lacking expression of the CAR target antigen. In three independent experiments, surviving CAR-T-treated mice underwent sequential CD19KO tumor rechallenges at escalating doses (1×10^5^, 1×10^6^, and 3×10^6^ cells per mouse). Age-matched naïve BALB/c mice receiving identical CD19KO tumor inocula served as untreated controls for the rechallenge. Survival data from all three experiments were pooled for analysis (n=25 CAR-T treated mice; n=30 untreated controls) and compared using the Kaplan-Meier method, with statistical significance determined by the log-rank test.

**In-vitro cytotoxicity assay**

Strain specific CAR-T or mouse primary NK cells were serially diluted in triplicate in a 96-well plate. Target cells (A20-WT, A20-B2m-KO, or BL3750) were stained with CFSE (5 µM, Thermo Fisher **#C34554**) for 10 min at 37 °C, quenched with complete medium, washed, and added at 1×10^5^ cells/well to CAR-T or control wells. After 4–24 h co-culture, wells were acquired by flow cytometry (NxT, Acoustic Focusing Cytometer, thermofisher**)** with FSC/SSC corrected using target-only controls. CFSE+ viable cells were counted, and specific lysis was calculated using % lysis = (1 - (CFSE⁺ cells in co-culture / CFSE⁺ cells in target-only wells)) × 100.

**Single-cell RNA sequencing**

Sample preparation. Two single-cell RNA-seq experiments were performed on A20-WT and B2*m*-knockout (B2*m*-KO) tumors harvested from syngeneic BALB/c mice. For the B2*m* engraftment study, liver tumor nodules were collected near endpoint (n = 4 per group) and processed as fresh single-cell suspensions for standard 10x Genomics Chromium library preparation. For the CAR-T treatment study, tumors were excised five days after CAR-T cell therapy from four cohorts: WT-Control (n = 5), WT-CAR-T (n = 5), B2m-KO-Control (n = 3), and B2m-KO-CAR-T (n = 3), and processed using the Chromium GEM-X Flex fixation workflow. For both experiments, tumors were dissociated by mechanical passage through a 70-µm cell strainer in PBS supplemented with 2% FBS, followed by red blood cell lysis with ACK Cell Lysis Buffer (Thermo Scientific, A1049201). For the Flex experiment, dead cells were removed using the EasySep Annexin V Dead Cell Removal Kit (STEMCELL Technologies, 17899) to achieve >90% viability. Single cells were then fixed using the Chromium GEM-X Flex Sample Preparation Kit v2 (10x Genomics) according to the manufacturer's instructions: cells were resuspended in Fixation Buffer B and incubated for 16–24 h at 4°C, quenched with Quenching Buffer B, and supplemented with Enhancer and glycerol to a final concentration of 10%. Fixed samples were stored at −80°C until library preparation. Cell concentration and viability were confirmed using a Nexcelom Cellometer K2 with ViaStain AOPI Staining Solution (Revvity).

Library preparation and sequencing. Gene-expression libraries were generated using either the standard 10x Genomics Chromium kit (experiment 1, targeting ~6,000 cells per sample) or the Chromium GEM-X Flex Gene Expression Kit v2 (experiment 2, targeting ~20,000 cells per sample). Cells were encapsulated into nanoliter gel beads-in-emulsion (GEMs) containing unique barcodes, and libraries were sequenced on an Illumina NovaSeq X at approximately 50,000 reads per cell.

Data processing. Sequencing reads were processed using Cell Ranger (v7.0.0, 10x Genomics) aligned to the mm10-2020-A mouse reference genome. Flex samples were processed with Cell Ranger multi using the fixed RNA-profiling workflow and the Chromium Mouse Transcriptome Probe Set (v1.0) for probe-based cell-barcode assignment and UMI quantification. Sixteen samples spanning the four experimental groups were processed separately and integrated for downstream analysis.

Quality control, integration, and clustering. Quality control and downstream analysis were performed using Seurat (v5.0.0 and v5.1.0) in R (v4.3.0). Cells were filtered using chemistry-appropriate thresholds: for the standard experiment, cells with <250 expressed genes, <500 UMIs, or >20% mitochondrial reads were excluded, and doublets identified by scDblFinder were removed; for the Flex experiment, cells with <500 detected genes, >8,000 detected genes, <1,000 UMIs, or >10% mitochondrial transcripts were excluded. Data were normalized using SCTransform with regularization, and batch effects between samples were corrected using Harmony on the first 30–50 principal components. Graph-based clustering was performed using the Louvain algorithm on a shared nearest neighbor graph (k = 20, resolution = 0.6), with visualization via UMAP.

Cell-type annotation. Cell types were annotated using SingleR with either the Mouse RNA-seq reference (MouseRNAseqData, Celldex) or the ImmGen reference dataset, validated by expression of canonical lineage markers (Cd3e/Cd3d for T cells, Cd79a/Ms4a1 for B cells, Cd14/Lyz2 for monocytes, Nkg7/Gzma for NK cells). CD4 and CD8 T-cell populations were re-clustered to identify transcriptionally distinct subsets.

Differential expression and pathway analysis. Differential expression between conditions was performed using limma and edgeR incorporating cellular detection rate as a covariate (experiment 1; adjusted p < 0.05, |log2FC| > 0.5), or using Seurat's FindMarkers function (min.pct = 0.05) with pseudo-bulk differential expression via Libra (v1.7) (experiment 2). Functional pathway enrichment was evaluated by Gene Set Enrichment Analysis (GSEA) using Hallmark gene sets from MSigDB.

Cell-cell communication. Cell–cell communication analysis was performed using MultiNicheNet (v2.1.0) with the NicheNet v2 mouse ligand–receptor network and ligand–target prior knowledge model. The analysis was restricted to the two CAR-T–treated groups (WT_CART, n = 5; B2m_CART, n = 3). Gene counts were aggregated into pseudo bulk profiles for each mouse across ten cell populations: CAR-T CD4, CAR-T CD8, eCD4, eCD8, monocytes, macrophages, neutrophils, NK cells, tumor B cells, and non-tumor B cells. Differential expression between WT_CART and B2m_CART was assessed using edgeR. MultiNicheNet predictions were manually curated to remove intracullular interactions not known to mediate cell-cell communication.

**IFNγ ELISPOT**

Human IFNγ ELISPOT

For quantifying the IFNγ response of endogenous (non-CAR) T cells from LBCL patients against tumors, an enzyme-linked immunospot (ELISPOT) assay was performed following manufacturer recommendations. On day 1, tumor samples were thawed, and CD3+ T cells were magnetically depleted using human CD3 MicroBeads (Miltenyi, #130-050-101). T cell-depleted-tumors were activated with 800ng/mL of recombinant human CD40L (Biolegend, #591706) and 2ng/mL of recombinant human IL-4 (R&D SystemS #204-IL-050) in complete RPMI at 37C and 5%CO_2_ for 2 days, as previously described^15,16^. The depleted CD3+ tumor infiltrating lymphocytes (TILs) were cultured in complete RPMI in a humidified incubator at 37C 5%CO_2_. and used as baseline (BL) samples for those patients that didn’t have other pre-CAR-T samples available. On Day 2, BL (PBMCs, aph PBMCs or TILs) and post-CAR PBMCs were thawed and CD19+ cells were magnetically depleted (Miltenyi, #130-050-301). CAR+ T cells were depleted from post-CAR samples using an PE-labelled human CD19 recombinant protein (Acro, #CD9-HP254) that specifically binds to the CAR, and PE microbeads (Miltenyi, #130-048-801). Both effector samples were cultured in complete RPMI in a humidified incubator at 37C and 5%CO_2_. Human IFNγ ELISPOT plates (BD Bioscience, #551849) were coated following manufacturer recommendations. On day 3, all cells were harvested and counted, and BL and post-CAR samples were co-cultured on IFNγ ELISPOT plates with autologous T-depleted tumor cells (tumor) or with 3T3-null or 3T3-CD19 cells as controls. Effector cells alone (unstimulated), effector cells stimulated with PMA and Ionomycin (PMA/Iono), and tumor and 3T3 cells alone were included as controls. IFNγ spots were developed after 48h following manufacturer instructions, and plates were scanned using CTL ImmunoSpot 3.0 reader. Samples were run in triplicates

Mouse IFNγ ELISPOT

Effector eCD4 and eCD8 T cells isolated from control or CAR-T-treated mice were harvested, counted and 5x10^4^ cells were cultured per well in a pre-coated IFNγ ELISPOT plate (Mabtech, #3321-4HPW-2) alone (unstimulated), stimulated with PMA/Iono, or co-cultured with the following target cells at 1:1 effector to target ratio (E:T): A20 WT, A20 CD19KO or A20 B2MKO (only for eCD8 cells). After 24h, IFNγ spots were developed following manufacturer instructions and plates were scanned using a Mabtech ELISPOT plate reader.

All samples were run in triplicates. Freshly made CAR-T cells were included as positive controls with the same conditions as for the eCD8 T cells. Each A20 cell type alone was included as negative control. Experiment was performed in three independent experiments with an n=3-4 mice per group each.

**Statistical Analysis**

Association of categorial variables with progression-free survival (PFS) and overall survival (OS) were performed in a univariable fashion, using Kaplan-Meier curves and a log-rank test. Association between signatures and complex genome features were implemented using Wilcoxon test. Statistical analysis of ELISPOTs were performed on GraphPad Prism using a Two-way ANOVA in all cases. For the patient ELISPOT a Tukey’s multiple comparison test was applied, while for the mouse ELISPOT, a Sidak’s or Dunnett’s (G, A20 WT vs B2MKO) multiple comparison test was used. A value of adjusted p<0.05 was considered significant.

**REFERENCES**

1	Nadeu, F. *et al.* IgCaller for reconstructing immunoglobulin gene rearrangements and oncogenic translocations from whole-genome sequencing in lymphoid neoplasms. *Nat Commun* **11**, 3390 (2020). <https://doi.org/10.1038/s41467-020-17095-7>

2	Rustad, E. H. *et al.* Revealing the impact of structural variants in multiple myeloma. *Blood Cancer Discov* **1**, 258-273 (2020). <https://doi.org/10.1158/2643-3230.BCD-20-0132>

3	Maura, F. *et al.* A practical guide for mutational signature analysis in hematological malignancies. *Nat Commun* **10**, 2969 (2019). <https://doi.org/10.1038/s41467-019-11037-8>

4	Rustad, E. H. *et al.* mmsig: a fitting approach to accurately identify somatic mutational signatures in hematological malignancies. *Commun Biol* **4**, 424 (2021). <https://doi.org/10.1038/s42003-021-01938-0>

5	Maura, F. *et al.* The mutagenic impact of melphalan in multiple myeloma. *Leukemia*, 1-6 (2021).

6	Jain, M. D. *et al.* Tumor interferon signaling and suppressive myeloid cells are associated with CAR T-cell failure in large B-cell lymphoma. *Blood* **137**, 2621-2633 (2021). <https://doi.org/10.1182/blood.2020007445>

7	Rustad, E. H. *et al.* Timing the initiation of multiple myeloma. *Nat Commun* **11**, 1917 (2020). <https://doi.org/10.1038/s41467-020-15740-9>

8	Consortium, I. T. P.-C. A. o. W. G. Pan-cancer analysis of whole genomes. *Nature* **578**, 82-93 (2020). <https://doi.org/10.1038/s41586-020-1969-6>

9	Shukla, S. A. *et al.* Comprehensive analysis of cancer-associated somatic mutations in class I HLA genes. *Nat Biotechnol* **33**, 1152-1158 (2015). <https://doi.org/10.1038/nbt.3344>

10	MacArthur, R. Species packing and competitive equilibrium for many species. *Theoretical Population Biology* **1**, 1-11 (1970). <https://doi.org/https://doi.org/10.1016/0040-5809(70)90039-0>

11	Kimmel, G. J., Locke, F. L. & Altrock, P. M. The roles of T cell competition and stochastic extinction events in chimeric antigen receptor T cell therapy. *Proc Biol Sci* **288**, 20210229 (2021). <https://doi.org/10.1098/rspb.2021.0229>

12	Gatenbee, C. D. *et al.* Immunosuppressive niche engineering at the onset of human colorectal cancer. *Nat Commun* **13**, 1798 (2022). <https://doi.org/10.1038/s41467-022-29027-8>

13	Berec, L. Impacts of Foraging Facilitation Among Predators on Predator-prey Dynamics. *Bulletin of Mathematical Biology* **72**, 94-121 (2010). <https://doi.org/https://doi.org/10.1007/s11538-009-9439-1>

14	McKay, M. D., Beckman, R. J. & Conover, W. J. A Comparison of Three Methods for Selecting Values of Input Variables in the Analysis of Output from a Computer Code. *Technometrics* **21**, 239-245 (1979). <https://doi.org/10.2307/1268522>

15	Malyguine, A. *et al.* A modified human ELISPOT assay to detect specific responses to primary tumor cell targets. *J Transl Med* **2**, 9 (2004). <https://doi.org/10.1186/1479-5876-2-9>

16	Neelapu, S. S. *et al.* Axicabtagene Ciloleucel CAR T-Cell Therapy in Refractory Large B-Cell Lymphoma. *N Engl J Med* **377**, 2531-2544 (2017). <https://doi.org/10.1056/NEJMoa1707447>
